# Does soil history decline in influencing the structure of bacterial communities of *Brassica napus* host plants across different growth stages?

**DOI:** 10.1101/2023.07.24.550396

**Authors:** Andrew J.C. Blakney, Marc St-Arnaud, Mohamed Hijri

## Abstract

Soil history has been shown to condition future plant-soil microbial communities up to a year after being established. However, previous experiments have also illustrated that mature, adult plants can “re-write”, or mask, different soil histories through host plant-soil microbial community feedbacks. This leaves a knowledge gap concerning how soil history influences bacterial community structure across different growth stages. Therefore, in this experiment we tested the hypothesis that previously established soil histories will decrease in influencing the structure of *Brassica napus* bacterial communities over the growing season. We used an on-going agricultural field experiment to establish three different soil histories, plots of monocrop canola (*B*. *napus*), or rotations of wheat-canola, or pea-barley-canola. During the following season, we repeatedly sampled the surrounding bulk soil, rhizosphere and roots of *B*. *napus* at different growth stages— the initial seeding conditions, seedling, rosette, bolting, and flower— from all three soil history plots. We compared the taxonomic composition and diversity of bacterial communities, as estimated using 16S rRNA metabarcoding, to identify any changes associated with soil history and growth stages on the different *B. napus* soil bacterial communities. We found that soil history remained significant across each growth stage in structuring the bulk soil and rhizosphere communities, but not the roots. This suggests that the host plant’s capacity to “re-write” different soil histories may be quite limited as key components that constitute the soil history’s identity remain present and continue to impact bacterial communities. For agriculture, this highlights how previously established soil histories persist and may have important long-term consequences on future plant-microbe communities, including bacteria.

## Introduction

Plants have never existed without microorganisms, hence the critical impact microbes have on plant metabolism, growth, and survival (1, 2, 3). For instance, soil microbes increase access to nutrients (4, 5, 6), temper environmental changes (7), or stress (8, 9), protect against pathogens (10, 11), and cue each plant developmental stage (7, 12, 13). For example, microbial communities can shift the timing or transition to different growth stages (GS) through nutritional or phytohormone pathways (13). The ubiquitous soil microbial communities help integrate these diverse signals and modulate the plant’s responses (9, 14). Such a long-term relationship, where plants interact with similar cohorts of microbial traits across generations, highlights a host plants capacity to tailor the structure of their bacterial communities in response to variable conditions and the plant’s needs through time (6, 15, 16, 17). As the soil is the source of the majority of plant-associated microbes (18), host plants have the capacity to alter the local soil chemistry through two concurrent processes; first, the host plant’s growth, development, and homeostasis, is determined by its capacity to uptake nutrients from the soil, which will influence the soil chemistry (19). Second, through rhizodeposition the host plant can vary the quantity and array of compounds released into the rhizosphere as required, which also changes the local soil chemistry (6, 15, 16, 17, 20).

As such, plant-soil microbial communities generate a reciprocal feedback process that incorporates various biotic and abiotic factors for the benefit of the current community (21, 22, 23), and which will impact future plant-soil microbial generations and their composition (24, 25, 26, 27). Thus, information from one plant-soil microbial community is transmitted through time to impact subsequent plant-microbial generations, i.e., that the soil history (SH), also referred to as soil legacy, of previous plant-soil microbial communities’ condition future ones (25, 28, 29). Future host plant can then alter the soil bacterial communities for their own purposes, in a phylogenetic-dependent manner (27, 30). However, several questions remain surrounding the duration of different SHs, what their impact on future plant-microbial communities may be, or how quickly plant-soil microbial community feedbacks may alter, or “re-write” different SHs.

Furthermore, SH will dictate how future plant-soil microbial communities form their microbial communities not only by establishing the biotic and abiotic context, but also through priority effects (31, 32, 33). As a new plant host develops, not only could its potential microbial community be constrained according to the SH, but also according to the composition at an earlier growth stage (GS). Unfortunately, however, most experiments testing SH have only focused on the microbial communities of adult plants (27, 29, 30, 34). Consequently, not incorporating temporal sampling has explicitly ignored the different roles of time on the assembly of microbial communities (35, 36, 37).

This has left a severe knowledge gap of how SH impacts microbial communities at different growth stages of the host plant, as well as how these communities develop throughout the growing season (38). For example, our previous experiment illustrated how different SHs could be “re-written” by host plant-soil microbial community feedback (PSF), or persist for up to a year (30, 34). However, since we only sampled adult host plants, we were unable to explore the strength or variation of different SHs on the microbial communities throughout the growing season. More recently, increased attention has been paid to seed (39, 40, 41, 42, 43) and flower microbial communities (44, 45, 46). These experiments focus on how microbes are vertically transmitted, subsequently establish new communities, and cue germination (13). In fact, microbes may be critical at early life stages; seed and seedling GS are precarious periods where the plant is most vulnerable to environmental stress and infection from phytopathogens (21, 38). Thus, experiments are needed to test the influence and dynamics of soil microbial communities at a variety of GS.

In this study, we investigated how different SHs impacted soil bacterial communities at five developmental GS throughout the season. We partnered with an on-going agricultural field experiment, as crop rotations and their agricultural inputs provide a ready-made model for how a previous PSF establishes a SH that impacts the future plant-bacterial community (22, 23). This helps address the lack of field experiments in studying SH (47). Here, the three established SHs were rotated plots of monocrop canola (*Brassica napus*), wheat-canola (WC), and pea-barley-canola (PBC). During the following season, we repeatedly sampled *B*. *napus* and surrounding soil at different GS— the initial seeding conditions, seedling, rosette, bolting, and flower—from all three SH plots. This design permitted us to test the hypothesis that the previously established soil histories—monocrop, WC, PBC, and their respective microbial communities and agricultural treatments—would decrease in influencing the structure of the *B*. *napus* bacterial rhizosphere communities over the growing season, as the host’s PSF reconstructs the existing soil history. To help us identify the competing influence of SH and GS, we compared the rhizosphere communities to their corresponding bulk soil (BS) and root communities at each GS. Bulk soil communities were a reference point to help account for the accumulative impact of time, the random ecological effects that can occur, and the different SHs on the soil bacterial communities throughout the growing season, but without the influence of the plant hosts at each growth stage. Conversely, the bacterial root communities were a reference point to account for the dominant influence of the host through time, but without the impact of the SH, as illustrated by our previous experiment (30).

We made four predictions that will help illustrate support for our “fading soil history through time” hypothesis: i) that the bacterial BS communities will remain stable, and continue to be primarily structured by their SH throughout the experiment, as they should not, by definition, be impacted by a plant host, ii) conversely, the bacterial root communities will be primarily structured by the different GS and shift accordingly, regardless of the SH, iii) the bacterial rhizosphere communities will be primarily structured by their SH at the seedling GS and resemble their cognate BS communities, and iv) as a result of the declining influence of SH, the bacterial rhizosphere communities will be primarily structured by PSF at the flower stage, and so will be more similar to one another and divergent from their cognate BS communities. To test our “fading SH through time” hypothesis, we estimated the bacterial communities from the BS, rhizosphere and RO of *B*. *napus* plants throughout the growing season using a 16S rRNA metabarcoding approach to infer amplicon sequence variants (ASVs). We compared the taxonomic composition and diversity of the bacterial communities to identify any changes associated with different SH and GS on the *B. napus* soil bacterial communities.

## Materials and Methods

### Site and experimental design

A long-term, crop rotation field experiment is on-going at the experimental farm of Agriculture and Agri-Food Canada’s Research and Development Centre, in Lacombe, Alberta (52°28′06″N, 113°44′13″W). The site is located in the semi-arid region of the Canadian Prairies; according to the weather station at the research farm, the 2019 growing season (May, June and July) had 197.8 mm of precipitation; compared to the 30-year average [1981-2010] of 216.3 mm. The daily temperature average for the 2019 season was 12.6 °C, while the 30-year average was 14.5°C. The farm has a loam texture (46% sand, 33% silt, and 21% clay), and has been well-described previously (48).

The experiment reported here was derived from the two-phase cropping sequence between the 2018 and 2019 growing seasons (Fig. S1A). The experimental design was a split-plot replicated in four complete blocks (Fig. S1B). For the 2018 Conditioning Phase, we selected three SH treatments that consisted of i) monocrop canola (*Brassica napus* L., cv. L252LL), ii) a two-year crop rotation between spring wheat (*Triticum aestivum* cv. AAC Brandon) and *B*. *napus* (WC) and iii) a three-year rotation between pea (*Pisum sativum* L. cv AAC Lacombe), barley (*Hordeum vulgare* cv. Canmore), and *B*. *napus* (PBC; Fig. S1). Thus, the 2018 Conditioning Phase established a SH composed of either canola, wheat, or barley, plus their respective management plans (30, 48). In the 2019 Test Phase, the 12 Conditioning Phase plots were all seeded with *B*. *napus*. The Test Phase established the *B*. *napus* host PSF, composed of the *B*. *napus* genotype, their soil bacterial community, their management plan, and previous SH (30, 34).

### Crop management and sampling

Crops were grown and maintained according to standard management practices, as previously described (48; see Supplementary Materials for details). Test phase *B*. *napus* plants were sampled at specific GS, seed, seedling, rosette, bolting, and flower, as described by the Canola Council of Canada (48). First, at the seed stage (GS00, May 10^th^, 2019), we took a 25 mL sample of the *B*. *napus* seeds to be seeded, as well an equivalent amount of soil from each plots. At the seedling stage, post-emergence, when only the cotyledons were visible (GS10, May 27^th^, 2019), five seedlings and their accompanying soil were pooled together for each sample. Composite samples were taken only at the seedling stage in order to have enough RO material for our subsequent DNA extractions. At the rosette, bolting and flower stages, individual plants and their associated soil were harvested from each plot. The rosette stage (GS19, June 18^th^, 2019) was harvested when nine leaves were visible, followed by the bolting stage (GS34, July 2^nd^, 2019) when a 20 cm stem was present. Finally, the flower stage (GS65, July 15^th^, 2019) was harvested when 50% of the flowers on the raceme were open (49).

At each growth stage we sampled three compartments: bulk soil (BS), rhizosphere, and root. Within each plot, BS was sampled from between the seeded rows, at least 10 cm from any seeds, or plants. Note that at the seed stage the only material collected was BS and the seeds. In the field, each plant had its aerial portions removed, and its roots and accompanying soil stored in coolers on ice. Based on the sampling strategy, in this study we define the BS microbiome as the soil bacterial community not influenced by the resident host plant, the rhizosphere microbiome as the bacterial community in the soil in close contact with the roots (29), and the root microbiome as the bacterial community attached to, and within, the roots (26). We accounted for the use of the various agricultural treatments in the downstream amplicon data by considering each plant sample and their total complement of particular agricultural treatments as a unit (30; see Supplementary Materials). In the field, all samples were kept on ice in coolers, then stored in the lab at -80°C before being shipped to Université de Montréal’s Biodiversity Centre, Montréal (QC, Canada) on dry ice for further processing (50, 51).

### DNA extraction from the Test Phase *B*. *napus* samples

Total DNA was extracted from all compartments (BS, seeds, rhizosphere, and roots) of the Test Phase field trial samples (see Supplementary Materials for details). Roots were first sieved out of the total soil sample and gently scraped it off using sterilized utensils into fresh collection trays. Seeds and root samples were ground separately in liquid nitrogen via sterile mortar and pestles (Fig. S2). For BS and rhizosphere samples, ∼500 mg was used for the NucleoSpin Soil gDNA Extraction Kit (Macherey-Nagel, Germany), while ∼130 mg of seeds and roots were used with the DNeasy Plant DNA Extraction Kit (Qiagen, Germany) (30, 34, 51). We failed to extract DNA from nine of the seedling root samples due to a lack of material; those samples were subsequently excluded hereafter (Fig. S2).

To estimate the composition of the bacterial communities in the BS, seed, rhizosphere and roots from the *B*. *napus* growth stages, extracted DNA from all samples were used to prepare 16S rRNA gene amplicon libraries using Illumina’s MiSeq platform (Génome Québec, Montréal) (30, 34, 51, 52). ASVs were then identified from the raw 11 010 728 MiSeq reads (53). We estimated the absolute abundance, or size, of the bacterial communities in each DNA sample by qPCR (30, 34, 54; see the Supplementary Materials for details).

### α-diversity of the Test Phase *B*. *napus* bacterial communities

In order to estimate the coverage of the bacterial domain of life, we calculated Faith’s PD as an α-diversity index from the *B*. *napus* samples using the pd function from the picante package (sum of all branch lengths separating taxa in a community; 55). To refine our understanding of the abundance and composition of the *B*. *napus* bacterial communities, we used two complementary methods to identify taxa specific to GS and SHs. First, taxa cluster maps were used to calculate the differential abundance of ASVs between experimental groups (56). Second, indicator species analysis was used to detect ASVs that were preferentially abundant in pre-defined environmental groups (compartments, GS, SHs; 57). Given the large number of taxa in our study, it was not practical to view taxa clusters as matrices below class, whereas indicator species analysis pinpoints specific ASVs of interest. Please refer to the Supplementary Materials for details.

### β-diversity of the Test Phase *B*. *napus* bacterial communities

To test for significant differences between the *B*. *napus* bacterial communities from different GS and SHs, we used the non-parametric permutational multivariate ANOVA (PERMANOVA). The PERMANOVA was calculated using the adonis function in the vegan package (58), with 9999 permutations, and the experimental blocks were included as “strata”. This was complemented with a PERMANOVA for each compartment (BS, rhizosphere and roots) that specifically tested GS and SHs as experimental factors, and used a weighted Unifrac distance matrix (59, 60) calculated using the distance function in phyloseq (61; see Supplementary Materials for details).

Distance-based redundancy analyses (RDA), using UniFrac distances weighted by absolute abundance, were used to quantified the amount of variation described by each experimental factors in the bacterial communities from the BS, rhizosphere, or roots (i.e. how much of the phylogenetic change between communities was due to the compartment, soil history, or GS; 57). Model accuracy was assessed with an adjusted *R^2^*value and tested for significance using an ANOVA (62).

## Results

Bulk soil and rhizosphere samples were more similar than to root or seed samples.

Illumina’s MiSeq produced 11 010 728 raw reads for the whole dataset, which were then processed through DADA2 (53, 63), where we retained 2 770 390 reads from all the experimental samples, which inferred a total of 33 392 ASVs (Table 1 & Fig. S3; see Supplementary Materials for details). Globally, we found that the bacterial communities from the BS, seeds, rhizosphere, roots were significantly different (PERM *R^2^* = 0.60135, *p* < 0.001). β-diversity analysis highlighted the difference between the root and seed communities from the BS and rhizosphere communities (Fig. S8A). These differences were further reflected by significantly different levels of phylogenetic diversity (PD), where the BS and rhizosphere communities remained the most diverse throughout the growing season (*p* < 0.001, Fig. S7). Comparatively, the roots and seed communities remained consistently less diverse (Fig. S7). Indicator species analysis did not identify any specific ASVs, according to compartment, growth stage (GS), nor soil history (SH).

**Table 1.** The bacterial bulk soil and rhizosphere communities had more unique ASVs than the root communities at each growth stage of their host plant *Brassica napus*. Samples were harvested throughout the 2019 growing season during the Test Phase of a multi-year crop rotation in Lacombe, Alberta. 11 010 728 raw reads were produced via Illumina’s MiSeq platform at Génome Québec, and processed through DADA2, where 2 770 390 reads were retained (16S rRNA Reads reported here) for ASV inference. A total of 33 392 ASVs were identified across the dataset. Bacterial community size was estimated by qPCR as the number of 16S rRNA gene copies.

| Growth Stage<sup>a</sup> | Compartment<sup>b</sup> | 16S rRNA Reads<sup>c</sup> | ASV Occurrence<sup>d</sup> | 16S rRNA Gene Copies<sup>e</sup> |
| --- | --- | --- | --- | --- |
| Seed | Bulk (n = 12) | 21 213 ±1842 | 1497 ±128 | 2 360 322 ±1 610 500 |
|  | Seed (n = 1) | 7 | 6 | 2 251 078 |
| Seedling | Bulk (n = 12) | 22 886 ±1516 | 1526 ±72 | 1 224 464 ±1 083 033 |
|  | Rhizosphere<br>(n = 12) | 19 817 ±3941 | 1355 ±143 | 6 722 767 ±1 929 973 |
|  | Root (n = 3) | 2519 ±117 | 432 ±28 | 1 153 665 ±738 663 |
| Rosette | Bulk (n = 12) | 23 020 ±4472 | 1448 ±128 | 1 415 797 ±1 088 160 |
|  | Rhizosphere<br>(n = 12) | 19 984 ±7475 | 1295 ±444 | 2 131 129 ±833 941 |
|  | Root (n = 12) | 3058 ±1099 | 361 S±67 | 13 105 921 ±5 757 254 |
| Bolting | Bulk (n = 12) | 26 811 ±6475 | 1567 ±209 | 3 266 621 ±1 896 126 |
|  | Rhizosphere<br>(n = 12) | 29 286 ±6090 | 1611 ±219 | 2 305 561 ±1 249 093 |
|  | Root (n = 12) | 2502 ±3362 | 231 ±116 | 8 711 539 ±4 380 029 |
| Flowering | Bulk (n = 12) | 28 704 ±6599 | 1649 ±150 | 3 049 801 ±3 738 808 |
|  | Rhizosphere<br>(n = 12) | 23 263 ±4147 | 1552 ±132 | 10 044 431 ±5 541 865 |
|  | Root (n = 12) | 4353 ±3784 | 352 ±105 | 3 159 861 ±1 327 309 |

a<sup>a</sup>, Test phase growth stages b<sup>b</sup>, Presented with the number of samples (n) retained c<sup>c</sup>, Values are presented as mean ± SD d<sup>d</sup>, Values are presented as mean ± SD e<sup>e</sup>, Values are presented as mean ± SD

### Bacterial bulk soil communities impacted more by soil history than growth stages

To test our hypothesis that the previously established SH would decrease in influencing the structure of the *B*. *napus* bacterial rhizosphere communities over the growing season, we first sought to establish the dynamics through time of the corresponding BS communities. We did this to help determine the impact of the different SHs through time on the bacterial communities, but without the influence of the plant-induced GS. Soil history and GS (i.e. time in the context of the bulk soil) were both significant in the BS communities, thought the interaction was not (PERM *R^2^* = 0.08770, *p* < 0.001; *R^2^*= 0.08596, *p* < 0.016, respectively; Table 2). PD remained stable across growing season, except at the flower stage where diversity was significantly higher (*p*. adj < 0.01; Fig. 1A). PD was also significantly higher at each GS time point in BS communities with a SH of PBC, compared to the communities from monocrop or WC plots (*p*. adj < 0.001; Fig. 1A). Monocrop BS communities were also globally depleted in *Fibrobacteria*, compared to WC or PBC bulk soils (*p*. < 0.05; Fig. 1B). The bacterial BS communities were enriched in class ABY1 (phylum *Patescibacteria*) at the rosette stage (*p*. < 0.05; Fig. S9B), and in class *Rhodothermia* (phylum *Bacteroidota*) at the bolting stage (*p*. < 0.05; Fig. S9C), when compared to their cognate rhizosphere communities.

**Figure 1.**
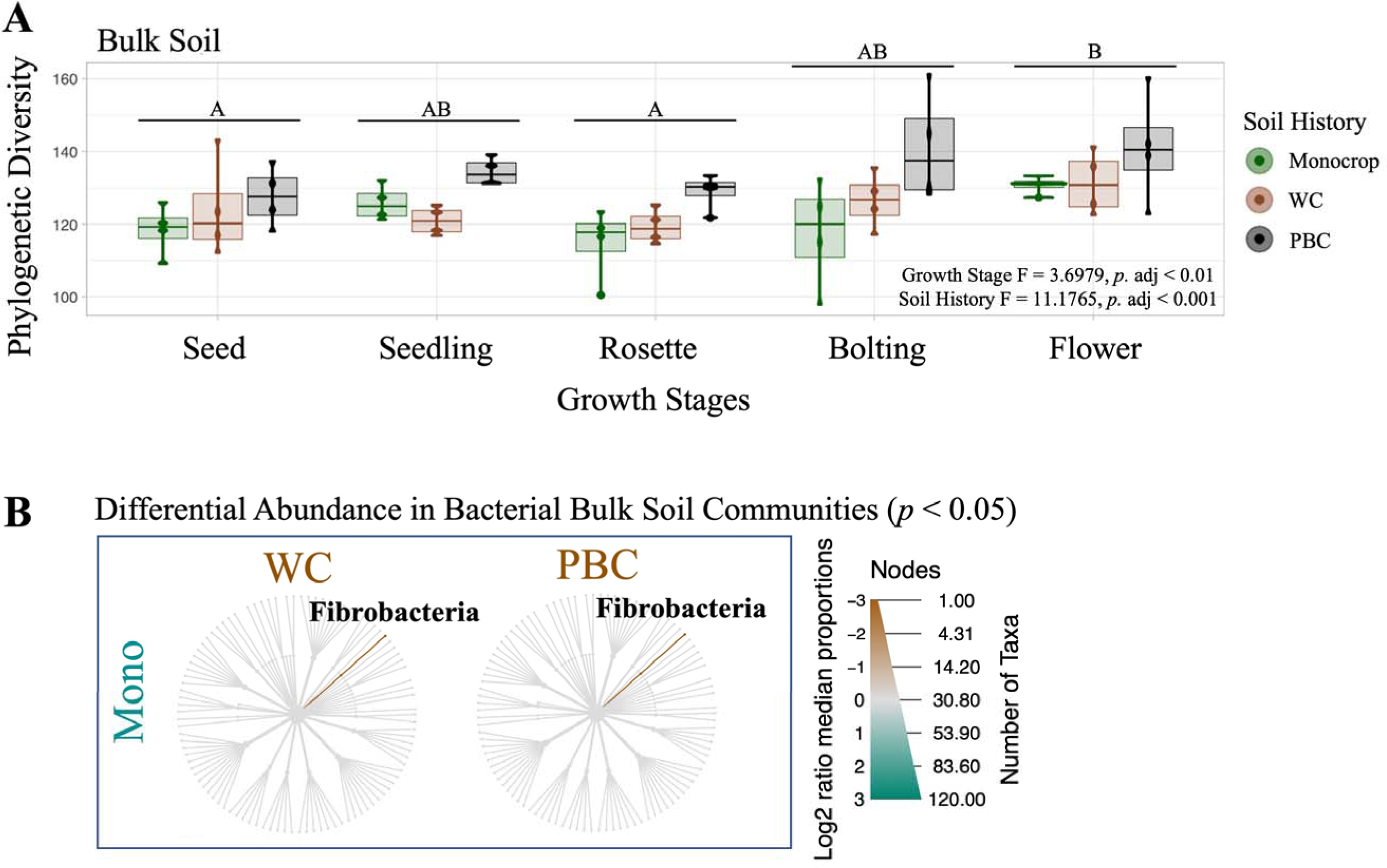
Bacterial communities identified from bulk soil samples were largely stable across different growth stages and soil histories. Samples were harvested throughout the 2019 *Brassica napus* growing season during the Test Phase of a multi-year crop rotation in Lacombe, Alberta. (A) Phylogenetic diversity was significantly higher during the flower growth stage than at the seed or rosette stages (A vs B, *p* < 0.01)). Bacterial bulk soil communities with the pea-barley-canola (PBC) soil history were also more diverse at each growth stage than the monocrop or wheat-canola (WC) communities (*p* < 0.001). Diversity across growth stages and soil histories was tested with a Multi-Factor ANOVA, which confirmed the previously established soil histories and the Test Phase *B*. *napus* growth stages did not interact. Statistically significant groups were identified using Tukey’s *post hoc* test. (B) *Fibrobacteria* were significantly enriched (*p* < 0.05) in bacterial bulk soil communities with wheat-canola (WC) and pea-barley-canola (PBC) soil histories, compared to communities from monocrop canola (Mono) plots. Significantly enriched taxa, labelled in bold, were tested between each pair of growth stages and soil history. Taxa that were significantly more abundant are highlighted brown or green, following the labels for each compared host. These differential taxa clusters identified significantly enriched (ie, abundant), using the non-parametric Kruskal test, followed by the *post hoc* pairwise Wilcox test, with an FDR correction.

**Table 2.** PERMANOVA identified growth stage and soil history as significant experimental factors for the bacterial bulk soil and rhizosphere communities harvested in 2019 from *Brassica napus* in the Test Phase of a multi-year crop rotation in Lacombe, Alberta. Soil history established the previous year was not significant for the bacterial communities from the root communities. PERMANOVA was calculated using a weighted Unifrac distance matrix, with 9999 permutations.

| Experimental<br>Factors | Test Phase Compartments<sup>a</sup> |  |  |  |  |  |  |  |  |
| --- | --- | --- | --- | --- | --- | --- | --- | --- | --- |
|  | Bulk Soil |  |  | Rhizosphere |  |  | Roots |  |  |
|  | F Model | R<sup>2</sup> | Pr (> F) | F Model | R<sup>2</sup> | Pr (> F) | F Model | R<sup>2</sup> | Pr (> F) |
| Growth Stage<sup>b</sup> | 1.31843 | 0.08596 | **0.016** | 2.04117 | 0.12043 | **0.001** | 2.33385 | 0.16454 | **0.006** |
| Soil History<sup>c</sup> | 2.69030 | 0.08770 | **0.001** | 2.73273 | 0.10749 | **0.001** | 1.40096 | 0.06585 | 0.163 |
| Soil History ~<br>Growth Stage | 0.71201 | 0.09285 | 0.928 | 0.70968 | 0.08374 | 0.812 | 0.95793 | 0.13508 | 0.548 |

<sup>a</sup>, Values in bold indicate significant factors or interactions
<sup>b</sup>, Test Phase growth stages: seed, seedling, rosette, bolting, or flower
<sup>c</sup>, Soil history established the previous year: monocrop canola (*B. napus*), wheat-canola rotation (WC), or pea-barley-canola rotation (PBC)

Redundancy analysis (RDA) also showed that the BS communities with PBC soil history were more phylogenetically consistent to each other across the growing season than to either the monocrop, or WC, as all the PBC soil communities remained more clustered (adj. *R^2^* = 0.0569, *p* < 0.001; Fig. 2A). We also observed that the BS communities at the seed and seedling communities tended to be more phylogenetically similar, while the remaining time points (rosette, bolting and flower) were more diverse (adj. *R^2^* = 0.0278, *p* = 0.009; Fig. 2B). Partitioning the β-diversity illustrated that the bacterial BS communities were dominated by turnover, or species replacement, throughout the growing season, regardless of their different SHs, though no transition between time points was significant (Fig. 2C).

**Figure 2.**
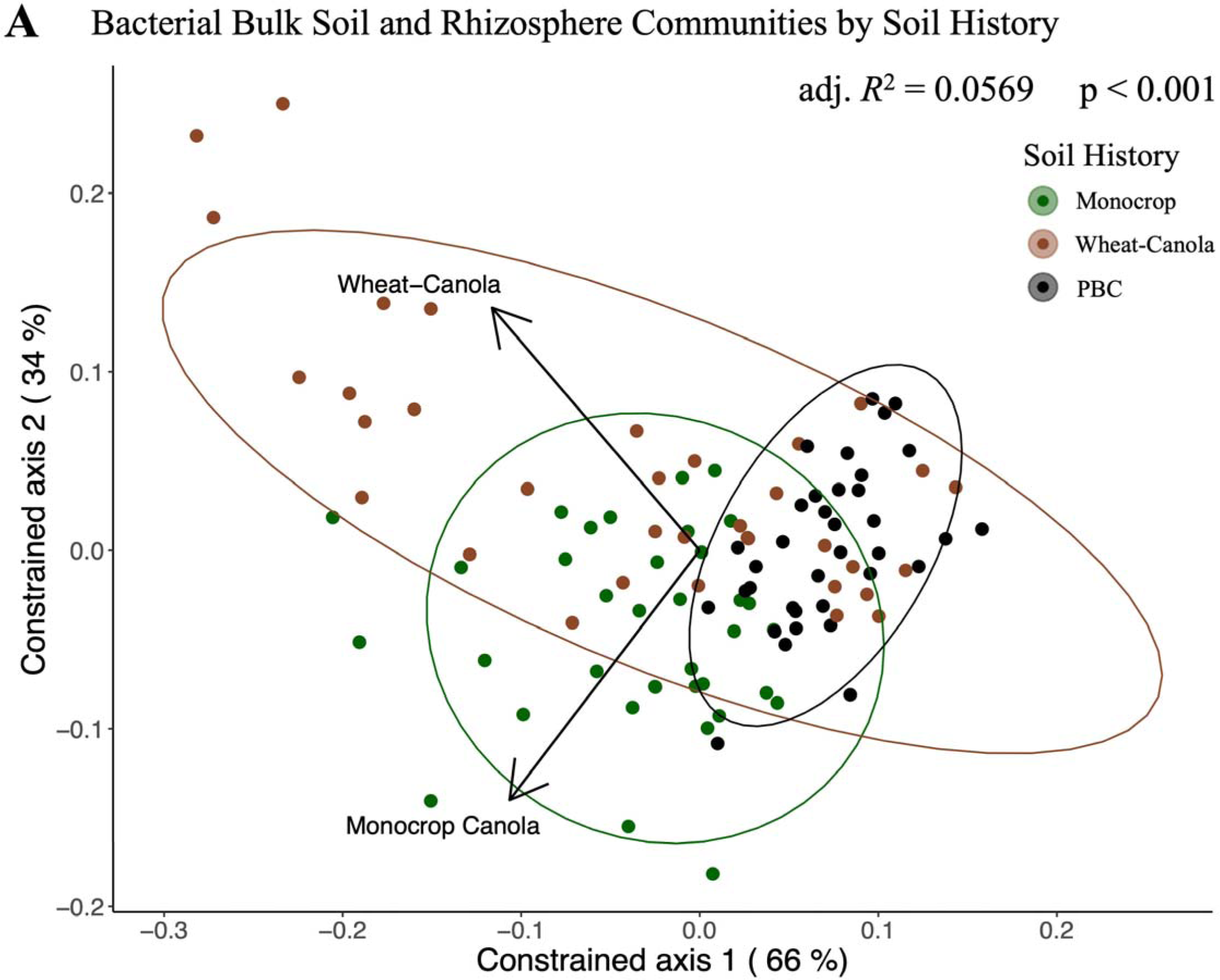
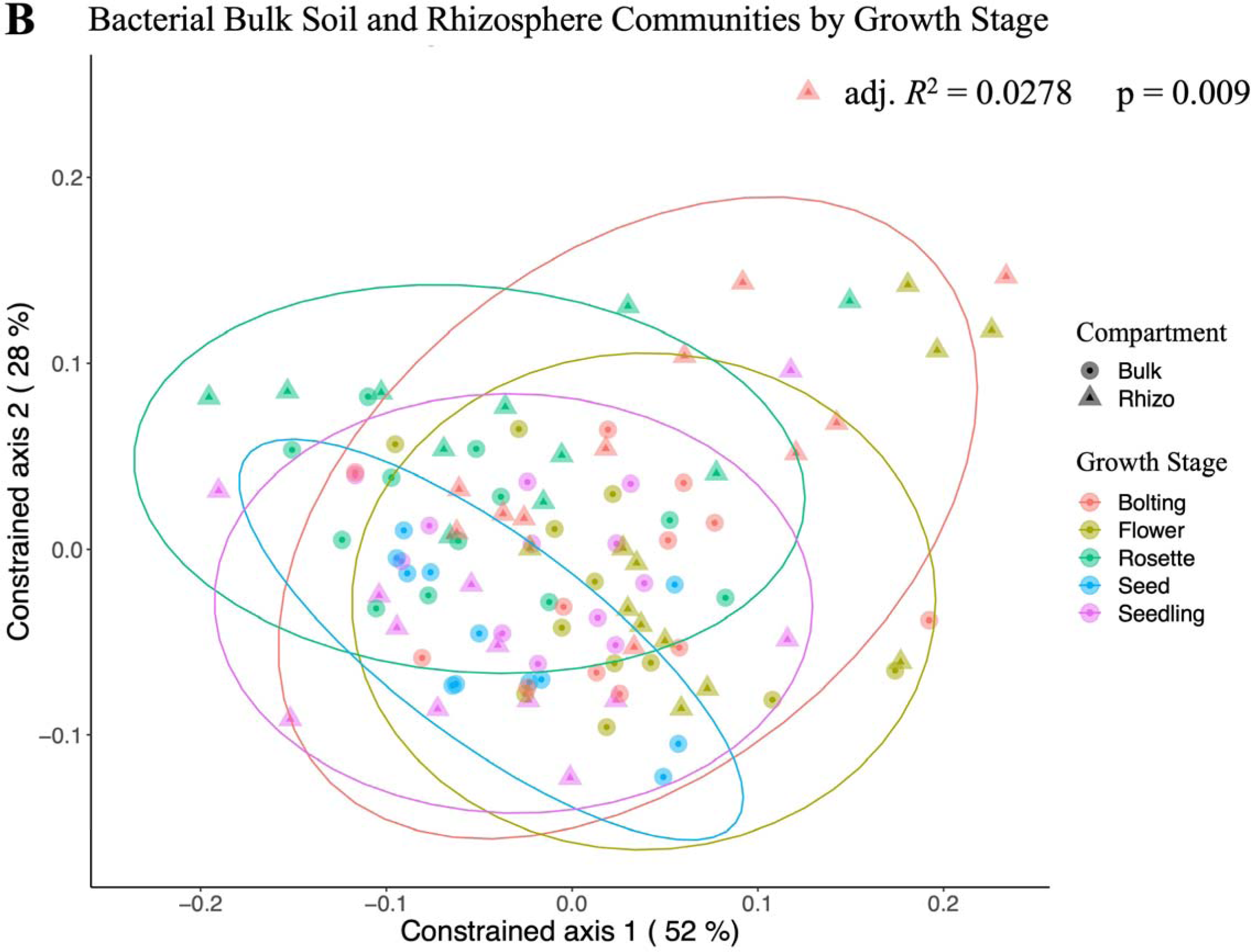
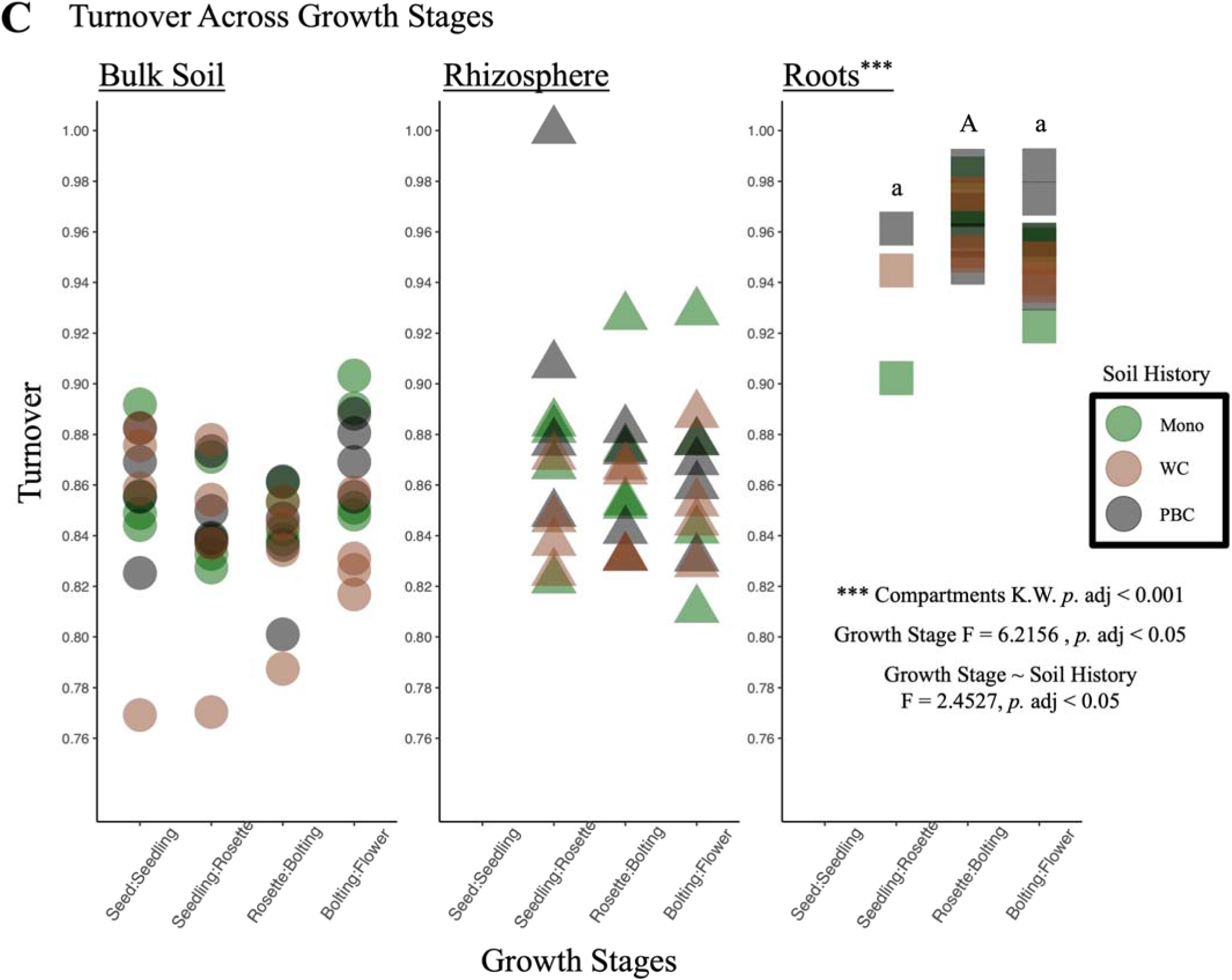
Bacterial communities identified from the bulk soil and rhizosphere of *Brassica napus* host plants were more phylogenetically similar in: (A) plots with pea-barley-canola soil history (black, PBC) versus other soil histories regardless of growth stage (adj. *R^2^* = 0.0569, *p* < 0.001), and (B) early growth stages (seed and seedling) versus later growth stages (rosette, bolting, and flower; adj. *R^2^* = 0.0278, *p* = 0.009). Samples were harvested throughout the 2019 growing season during the Test Phase of a multi-year crop rotation in Lacombe, Alberta. Distance-based redundancy analysis, using UniFrac distances weighted by absolute abundance, quantified how the experimental factors (A, constrained by soil history) and (B, constrained by growth stages) structured the bacterial communities, where those with similar phylogenetic composition appear closer together. (C) Beta-diversity of bacterial communities were dominated by turnover across each growth stage of *B*. *napus*, particularly in the roots (*p* < 0.001). Turnover was significantly higher in the roots during the transition from rosette to bolting (A vs a, *p*. adj < 0.05). Nestedness remained below 0.1 in all samples. Bray-Curtis dissimilarity distances from each communities within a plot were compared across each growth stage. To determine any significant differences in turnover among compartments the non-parametric Kruskal test was used, followed by the post-hoc pairwise Wilcox test with an FDR correction, as the data were not normally distributed. A Multi-Factor ANOVA confirmed that the turnover between growth stages, within each compartment, was only significant in the roots, and only interacted with soil history in the roots. Statistically significant groups were identified using Tukey’s post-hoc test.

### Bacterial root communities were only impacted by growth stage, and not soil history

Next, to establish the influence of the plant-induced GS, but with minimal impact from the different SHs, we analyzed the dynamics of the bacterial root communities across the different GS. Here we found that only GS was significant in structuring root communities (PERM *R^2^* = 0.16454, *p* < 0.006; Table 2), as opposed to the BS communities where both the time points and SH were significant. RDA further illustrated the significant impact of GS on the roots, with no impact of SH (adj. *R^2^* = 0.0569, *p* < 0.001; Fig. 3C). We also observed that the root communities were more phylogenetically consistent at the seedling stage and became more variable at each subsequent GS (Fig. 3C). Finally, partitioning the β-diversity also highlighted the importance of GS in the roots, as turnover was significantly higher during transition from rosette to bolting in the roots (*p.* adj < 0.05; Fig. 2C). Bacterial community turnover was also significantly higher in the roots than in the BS, or rhizosphere (*p*. adj < 0.001; Fig. 2C).

**Figure 3.**
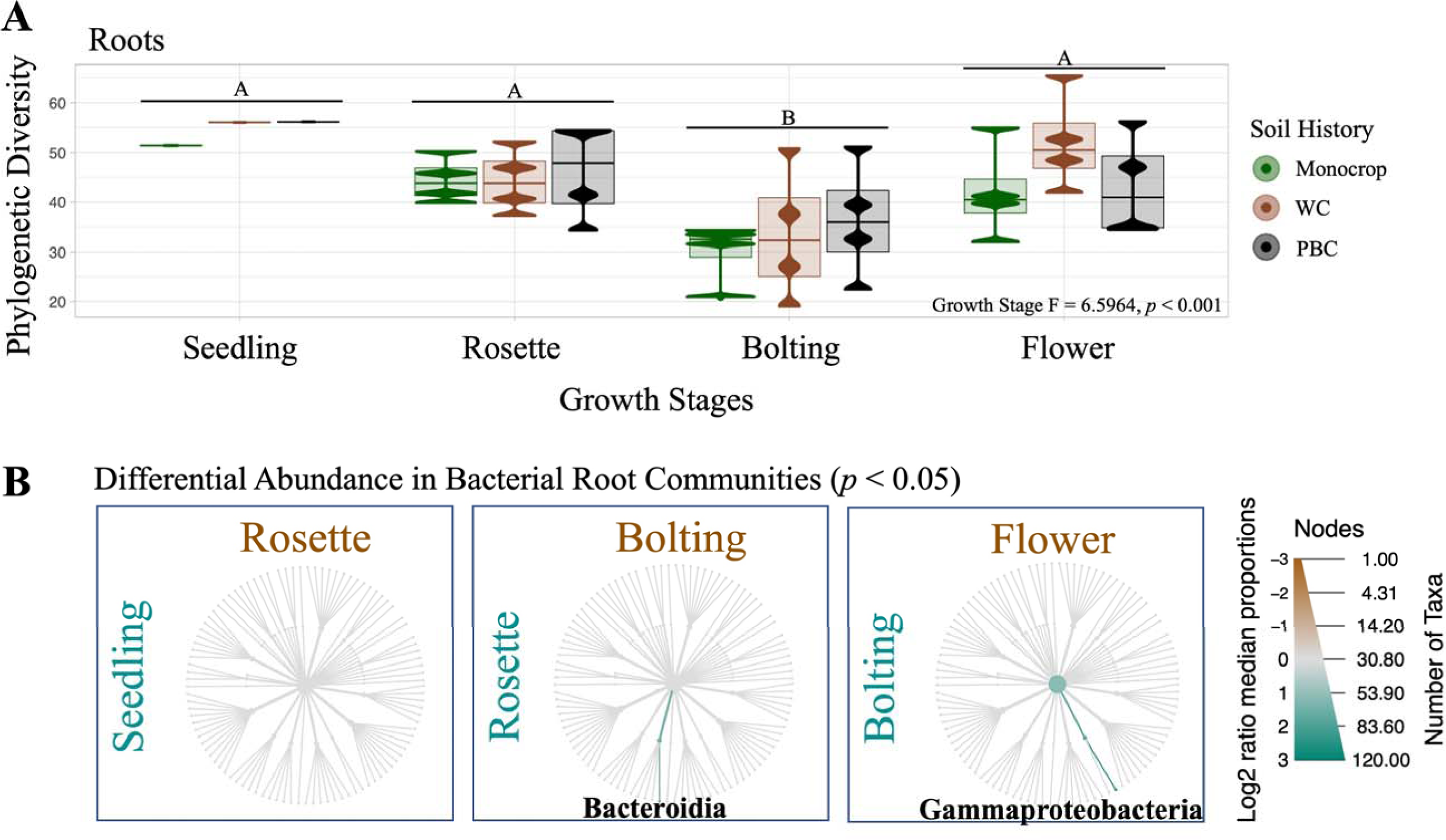
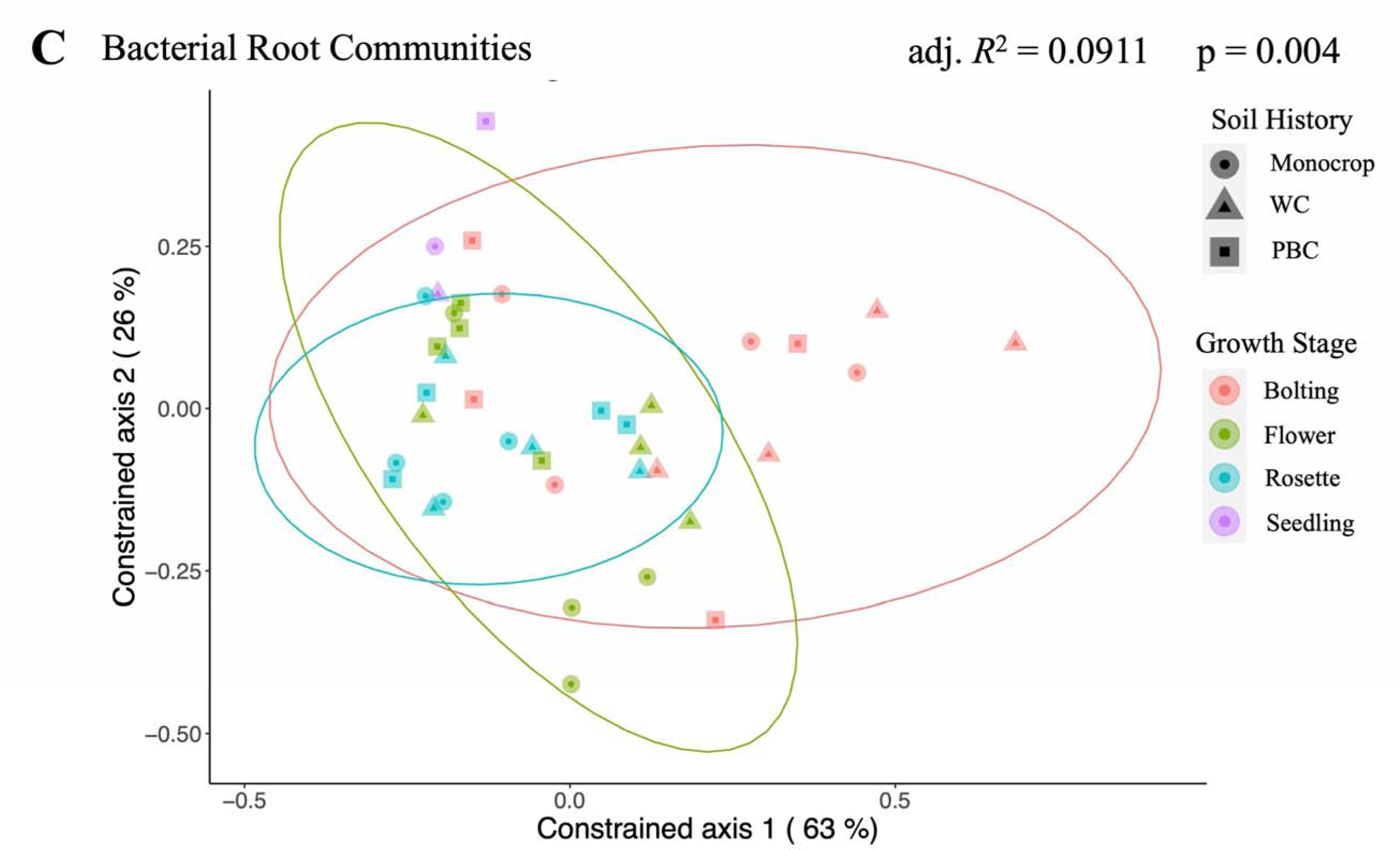
Bacterial root communities were stable across different growth stages and soil histories. Samples were harvested throughout the 2019 *Brassica napus* growing season during the Test Phase of a multi-year crop rotation in Lacombe, Alberta. (A) Phylogenetic diversity was significantly lower among the root communities at the bolting stage compared to other growth stages (*p* < 0.001). Diversity across growth stages was tested with a Multi-Factor ANOVA, which confirmed that the previously established soil histories were not significant and did not interact with the Test Phase *B*. *napus* growth stages. Statistically significant groups were identified using Tukey’s *post hoc* test. (B) *Bacteroidia* were significantly enriched (*p* < 0.05) in the bacterial root communities at the rosette stage compared to the subsequent bolting stage, while *Gammaproteobacteria* were enriched in root communities at the bolting stage compared to communities at the flower stage. Significantly enriched taxa, labelled in bold, were tested between each pair of growth stages and soil history. Taxa that were significantly more abundant are highlighted brown or green, following the labels for each compared host. These differential taxa clusters identified significantly enriched (ie, abundant), using the non-parametric Kruskal test, followed by the *post hoc* pairwise Wilcox test, with an FDR correction. (C) Bacterial root communities from the seedling (purple) and rosette stage (blue) were more phylogenetically similar to each other, regardless of different soil histories (shapes; not significant), than to the bolting (red) or flower (green) growth stages (adj. *R^2^*= 0.0911, *p* = 0.004). Distance-based redundancy analysis (constrained by growth stages), using UniFrac distances weighted by absolute abundance, quantified how the experimental factors structured the bacterial communities, where those with similar phylogenetic composition appear closer together.

In the root communities, PD was low compared to the BS and rhizosphere communities (Fig. S7). We did not detect any impact of the different SHs on the PD, nor differential abundances of taxa, in the root communities at any GS. These communities were very stable across GS, except at the bolting stage, where PD was significantly lower compared to the other GS (*p*. adj < 0.01; Fig. 3A). The root communities were significantly depleted in a wide variety of bacterial taxa at each GS when compared to the rhizosphere (*p* < 0.05; Fig. S9C). Nevertheless, the roots were also significantly enriched in specific taxa, compared to the rhizosphere (Fig. S9). First, at the rosette stage, the most prominently enriched taxa in the roots were in the *Verrucomicrobiae*, *Actinobacteria*, *Proteobacteria*, and *Bacterodia* (*p* < 0.05; Fig. S9B). Second, at the bolting stage, root communities were enriched in the *Gammaproteobacteria* & *Bacterodia* (*p* < 0.05; Fig. S9C). We also detected two enriched taxa in the roots when compared among themselves at different GS; *Bacterodia* were enriched at the rosette stage, compared to the bolting stage, while *Gammaproteobacteria* were enriched at the bolting stage, compared to the flower stage (*p* < 0.05; Fig. 3B).

### Rhizosphere communities were impacted by growth stages and soil history

Finally, we compared how the dynamics of the bacterial rhizosphere communities were influenced by GS and SH. Similar to BS communities, SH and GS were also both significant for the rhizosphere communities, while the interaction was not (PERM *R^2^* = 0.10749, *p* < 0.001; *R^2^* = 0.12043, *p* < 0.016, respectively; Table 2). RDA illustrated that the rhizosphere communities remained phylogenetically similar with the BS communities across the growing season, while being most similar at the seedling stage and the initial seed stage in the BS (Fig. 2B). Also like the BS communities the PBC communities were more phylogenetically consistent to each other, regardless of GS, than to either the monocrop, or WC (Fig. 2A).

Rhizosphere communities with PBC soil history also had significantly more PD at each GS, than the rhizosphere communities from either the monocrop or WC soil histories (*p*. adj < 0.001; Fig. 4A). However, PD generally increased across GS regardless of previous SH, such that the bolting and flower rhizosphere communities were more diverse than those from the seedling and rosette GS (*p*. adj < 0.001; Fig. 4A). A similar trend was observed in the β-diversity, such that rhizosphere communities had more variation at the rosette, bolting and flower stages compared to communities in the BS (Fig. 2B). The β-diversity of the rhizosphere communities was also driven by turnover across each GS, and was unaffected by different SH, similar to the BS communities (Fig. 2C).

**Figure 4.**
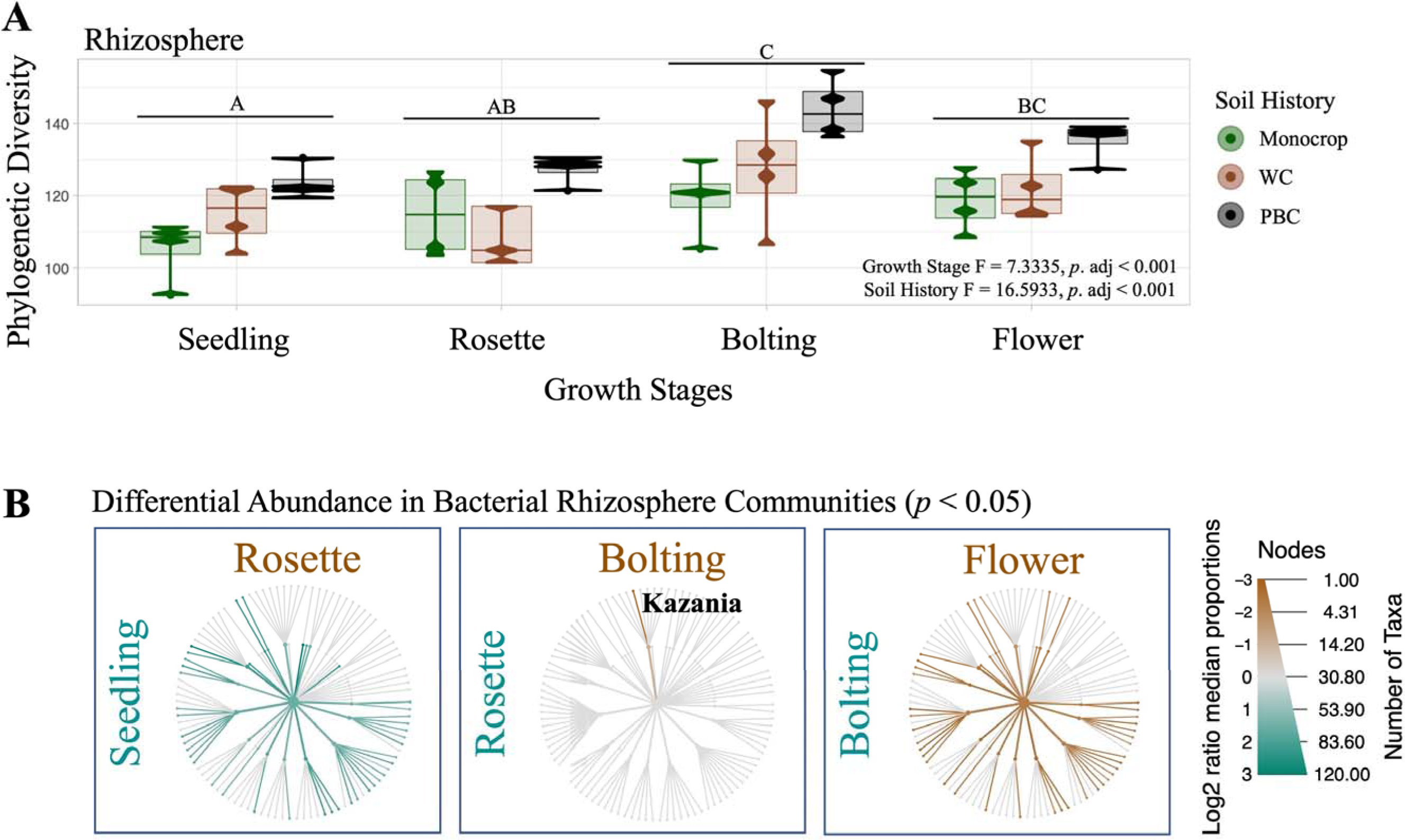
Bacterial rhizosphere communities varied significantly between growth stages of their *Brassica napus* plant host. Samples were harvested throughout the 2019 *Brassica napus* growing season during the Test Phase of a multi-year crop rotation in Lacombe, Alberta. (A) Phylogenetic diversity was significantly higher in the bolting and flower growth stages than the seedling and rosette stages (*p* < 0.001). Bacterial rhizosphere communities with the pea-barley-canola (PBC) soil history were also significantly more diverse at each growth stage than the other soil histories (*p* < 0.001). Diversity across growth stages and soil histories was tested with a Multi-Factor ANOVA, which confirmed the previously established soil histories and the Test Phase *B*. *napus* growth stages did not interact. Statistically significant groups were identified using Tukey’s *post hoc* test. (B) The seedling-rosette transition and bolting-flower transition were significantly different (*p* < 0.05) in bacterial taxa. Significantly enriched taxa, labelled in bold, were tested between each pair of growth stages and soil history. Taxa that were significantly more abundant are highlighted brown or green, following the labels for each compared host. These differential taxa clusters identified significantly enriched (ie, abundant), using the non-parametric Kruskal test, followed by the *post hoc* pairwise Wilcox test, with an FDR correction.

We observed a number of significant changes to taxonomic abundances in the rhizospherre, compared to the BS and root communities, at several GS (*p* < 0.05; Fig. S9), with the most prominent and widespread enrichment being during the seedling and flower stages (*p* < 0.05; Fig. 4B). The rhizosphere communities were largely stable amongst themselves between the rosette and bolting stages in terms of different taxa abundance, except for an enrichment in *Kazania* bacteria (phylum *Desulfobacterota*; Fig. 4B). However, they were noticeably enriched in a number of taxa compared to the RO at the rosette and bolting stages (Fig. S9B & C). There were no ASVs that were significantly enriched, or depleted, in the rhizosphere according to GS or SH.

## Discussion

Soil history implies that information is transmitted through time to condition the assembly of future plant-soil microbial communities (25, 28, 29). Although our previous work has shown this can be true (30, 34), the influence or longevity of SH on bacterial community structure across growth stages is not clear. In this study, we tested how previously established SH endured across GS throughout a growing season. We hypothesised that previously established SHs will decrease in influencing the structure of *Brassica napus* bacterial rhizosphere communities over the growing season. We took advantage of an agricultural field experiment to bridge the gap between controlled greenhouse conditions and experiments in “natural” environments, as such studies are currently lacking to understand bacterial temporal dynamics (47, 64, 65). We sampled bulk soil, rhizosphere and roots successively throughout the *B*. *napus* growing season from plots with different SHs and used 16S rRNA metabarcoding to identify the bacterial communities. Contrary to our hypothesis, the weight of evidence suggests that the different soil histories did not decline in influence, but rather continued to have a significant impact on the bacterial rhizosphere communities throughout the growing season.

These communities are initially established at the confluence of the soil environment and the immediate space the plant host may be able to influence (29, 66). Given that the new *B*. *napus* host would be mere days emerging from its seed into a new soil environment with an established SH (66), and minimally able to exert much influence on the surrounding soil through PSF (38), we predicted that at the early seedling GS the bacterial rhizosphere communities would be primarily structured by their SHs and resemble their cognate BS communities. Indeed, we found that the bacterial communities at the seed and seedling time points from the BS and rhizosphere were more phylogenetically similar than at later time points, or between different SHs (Figs. 1, 2, & 4). Nonetheless, we also observed that within this phylogenetic similarity there were widespread taxonomic enrichments in the rhizosphere at the seedling GS, compared to the BS and root communities (Fig. S9). This suggests that at initial GS the SH constrains the composition of the bacterial communities, while individual taxa may be attracted to the new rhizosphere from the surrounding soil and enriched. Future work is needed to determine how active the seedling is in this early attraction.

Across the subsequent GS, the most striking observation were the widespread community dynamics and taxonomic enrichments in the bacterial rhizosphere communities (Fig. 4B & S9). This was entirely absent in the BS communities (Fig. 1B & S9) and more limited in the roots (Fig. 3 & S9), although GS was significant in structuring all three communities (Table 2). The root communities should be primarily shaped by the plant host across GS, as we predicted and observed (Table 2). Conversely, the BS communities should largely experience similar abiotic conditions as the rhizosphere, but with minimal influence from the plant host (66). As such, we predicted that these communities would remain largely stable through time, though we actually found that the BS communities were also significantly structured at each time point (Table 2). The significance of the time points structuring the BS communities is likely a reflection of the aging soil.

In fact, the increased variation and phylogenetic diversity was a key feature we detected in both the rhizosphere (Fig. 2B & 4) and BS communities (Fig. 1A). We predicted that due to the declining influence of SH, the bacterial rhizosphere communities would be primarily structured by PSF through time. Therefore, we expected that by the flower GS the rhizosphere communities would be more similar to one another, regardless of their SH, and more divergent from their cognate BS communities. We did confirm that the rhizosphere communities varied significantly from their BS communities (Fig. 2B & S9), and we observed an increase in PD at later GS (Fig. 1A & 4A), which both endorses the prediction. It also mirrors the root communities, which were also significantly influenced by GS, though never by SH (Table 2), and also demonstrated increasing variation through time (Fig. 3). In both the BS and rhizosphere communities we observed increased PD over time, as older soils can slowly accumulate new community members due to different dispersal, drift, selection, or speciation/diversification events (67).

However, rhizosphere communities are also affected by the changing influences of the *B*. *napus* plant host at each GS, while those in the BS are not since they are non-planted controls. Thus, we ought to consider that the trend among the rhizosphere communities may also be due in part to the plant host, unlike in the BS. Similar increases in PD across GS have also been observed in the rhizosphere of other plants, such as rice (68). Unlike previous findings in the perennial *Brassicaceae Arabidopsis alpina* that showed quite static bacterial communities (69), our data aligned similarly to other annual crops that exhibit dynamic bacterial rhizosphere communities across GS (68). This late GS variation observed in all three communities could also be due to the inherent stochasticity of priority effects, where community composition at later GS is constrained by earlier GS (31, 32, 33). This could be better tested in the future as high-frequency sampling from multiple hosts of the same genotype may be experimentally valid to detect priority effects (70).

Critically, we did not find that flower GS rhizosphere communities were more similar to one another, regardless of their SH, as expected (Fig. 2B). In our previous work we found that diverse *Brassicaceae* hosts at the flower GS converged toward phylogenetically similar rhizosphere communities, regardless of their different SHs (30). In our current experiment, the plant’s GS clearly do become more important in structuring the rhizosphere throughout the growing season. However, additional factors also continue to add diversity and prevent more convergence among the rhizosphere communities. This could further highlight the impact of priority effects in shaping subsequent bacterial communities (31, 32, 33). Alternatively, including other diverse host plants could better reveal how similar within-host rhizosphere communities are at different GS (30).

### Bacterial diversity was higher with more diverse soil history

Our results further illustrated that bacterial BS and rhizosphere communities with the PBC history consistently had the highest PD, compared to the monocrop and WC SH across all GS (Fig. 1A & 4A). We can be confident that the increased PD among the PBC soil communities was due to the different SH for two reasons. First, the increase in PD among PBC plots in the BS communities was present from the first GS, before any plant hosts was even present. Second, the increase in PD in the BS and rhizosphere communities from the PBC plots remained throughout the growing season. Thus, even with the addition of a plant host the common PBC soil history still impacted the bacterial communities at each GS, or concordant sampling time in the case of the BS.

Conversely, we found no difference between the three SHs in the root communities (Fig. 3), where SH was not significant to those communities (Table 2). Thus, from our experiment we can only observe a change in the root communities according to GS (Fig. 3). This is consistent with our previous work that also demonstrated root communities tend to be strongly influenced by the host plant, and weakly impacted by SH, unless the plant host is stressed, and unable to contribute to PSF (30, 34).

Although we found a clear impact of different SHs on PD in the bacterial BS and rhizosphere communities, it is interesting to note that we only identified a slight compositional change in the BS due to SH, but not the rhizosphere (Fig. 1B). Moreover, we did not detect specific ASVs within rhizosphere communities according to their different SHs. This lack of compositional difference between SH might suggest that the different agricultural treatments involved in establishing the previous SH were not sufficiently diverse. However, given our previous results using crop rotations this seems unlikely (30, 34). Alternatively, the lack of compositional differences between rhizosphere communities despite coming from different SHs, could be evidence for the common host plant structuring similar rhizosphere communities. This would be consistent with other studies that found declining site-specific bacteria over time, and increasing plant-specific bacteria. Including other diverse host plants, similar to our previous experimental design (30, 34), would allow us to better support this conclusion.

## Conclusion

In this experiment we tested the hypothesis that previously established soil histories would decrease in influencing the structure of *Brassica napus* bacterial rhizosphere communities over the growing season. We largely confirmed our first and second predictions, which had suggested that the bacterial bulk soil and root communities would be primarily structured by soil history and growth stages, respectively. Our results for rhizosphere communities at the initial seed and seedling stages also confirmed our prediction that these communities would remain similar to their corresponding bulk soil communities and soil histories. We also found that the bacterial bulk soil, rhizosphere, and root communities all diverged more as they aged. However, the rhizosphere communities did not converge in similarity over the growing season, regardless of their soil history, refuting our final prediction. In fact, soil history continued to be influential among the rhizosphere communities across the different growth stages. Furthermore, we show that soil histories established with more plant diversity contribute to more phylogenetically diverse bacterial communities. Therefore, based on our data, our initial hypothesis concerning the decline of soil history influencing the structure of the bacterial rhizosphere communities across the growing season was only partly supported. Instead, we found a strong impact of soil history on the bacterial rhizosphere communities throughout the growing season that stressed the nuanced interactions of plant-soil microbial community feedback.

Our results highlight the importance of studying microbial communities through time, which has largely been ignored to date. Studying how communities arrive at a given composition is more instructive than just a static snapshot. Here we found that different soil histories persisted and impacted bacterial diversity throughout the growing season. This suggests that the host plant’s capacity to “re-write” different soil histories may be quite limited as key components that constitute the soil history’s identity remained present and continued to impact the bacterial communities. From the agricultural perspective, persisting soil histories may have important long-term consequences, such as building capacity for more resilient, diverse bacterial communities (22). This presents exciting future experiments to uncover the transmission components, or memory, of soil history among soil bacterial communities. Given the significant and myriad human-induced changes throughout the biosphere (71), there is a clear need to better account for how historical events may structure plant-soil microbial communities going forward through time, and more broadly influence the mechanisms of community ecology.

## Supporting information

Blakney et al., 2023 BioRxiv Supplemental Materials

## Acknowledgements

This work was supported by the Natural Sciences and Engineering Research Council of Canada to MH (Grant Numbers: CRDPJ 500507-16 and RGPIN-2018-04178) and Canola Council of Canada, which are gratefully acknowledged. We thank Chantal Hamel for her support and assistance in establishing the field experiment with Agriculture and Agri-food Canada. We also thank Jennifer Zuidof for setting up, managing, and harvesting the field experiment at Lacombe, Albert. We thank Stéphane Daigle for assistance in statistical analysis, Jacynthe Masse, Simon Morvan, Morgan Botrel, and Stephanie Shosha, for their helpful comments and discussions.

## Author Contributions

AJCB designed the experiment, prepared the samples for sequencing, performed the qPCR experiment, analyzed the data, and wrote the manuscript with input from all co-authors. MSA & MH designed the experiment, supervised the work, contributed reagents, analytical tools, and revised the manuscript.

## Data Accessibility

Sequencing data and metadata are available at NCBI Bioproject under accession number: PRJNA997731.

