## Supplementary material for "Does soil history decline in influencing the structure of bacterial communities of *Brassica napus* host plants across different growth stages?": Blakney et al., 2023 BioRxiv Supplemental Materials

List of Supplementary Tables

Table S1. Mock Community Composition
Table S2. Primers
Table S3. PERMANOVA

List of Supplementary Figures

Figure S1. A) Conceptual Design, & B) Experimental Design
Figure S2. Lab Workflow
Figure S3. DADA2 Workflow
Figure S4. Mock Community Results
Figure S5. Rarefaction Curve
Figure S6. 16S rRNA Standard Curve for qPCR
Figure S7. Phylogenetic Diversity of Compartments at Each Growth Stage
Figure S8. Principal Co-Ordinate Analysis of Weighted Unifrac Communities A) All Compartments, & B) Bulk soil and Rhizosphere Communities
Figure S9. Differential Abundance of Bacterial Taxa A) Seedling, B) Rosette, C) Bolting, & D) Flower

### Supplementary Methods

#### Crop management

A pre-seed ‘burn off’ herbicide treatment using glyphosate (Roundup, 900 g acid equivalent per hectare, a. e. $ha^{-1}$) and bromoxynil (Pardner, 280-330 g active ingredient per hectare, a.i. $ha^{-1}$) was applied to all plots each year to ensure a clean starting field prior to seeding. The herbicide Liberty was applied to *B. napus*, while Pixxaro A & B with Axial were applied to wheat and barley plots, for in-season weed control, while fungicides were only applied as needed. Soil tests were used to determine the rates of in-season nitrogen, phosphorus, and potassium application. Crops were harvested between late August and early October, depending on the crop and year. The subsequent Test Phase *B. napus* plant hosts were subjected to the same standard management practices as the Conditioning Phase, including pre-seed ‘burn off’, in-season herbicide and fungicide treatments as needed, and fertilized as recommended by soil tests.

#### DNA extraction from Test Phase *Brassicaceae* root and rhizosphere samples

For bulk soil and rhizosphere samples, ~500 mg was used for the NucleoSpin Soil gDNA Extraction Kit (Macherey-Nagel, Germany), while ~130 mg of seeds and roots were used with the DNeasy Plant DNA Extraction Kit (Qiagen, Germany) (Lay *et al*., 2018; Blakney *et al*., 2022). A no-template extraction negative control was used with the bulk soil, rhizosphere and root extractions, and included with the Test Phase samples (Fig. S2), to assess the influence of the extraction kits on our sequencing results, and the efficacy of our lab preparation. All extracted DNA samples were quantified using the Qubit dsDNA High Sensitivity Kit (Invitrogen, USA), and qualitatively evaluated by mixing ~2 µL of each sample with 1 µL of loading dye containing Gel Red (Biotium) and running it on a 0.7 % agarose gel for 30 minutes at 150 V. The no-template extraction negative controls were confirmed to not contain DNA after extraction, where the detection limit was > 0.1 ng (Qubit, Invitrogen, USA).

#### 16S rRNA gene amplicon generation and sequencing to estimate the bacterial community

First, all DNA samples were diluted 1:10 into 96-well plates using the Freedom EVO100 robot (Tecan, Switzerland). To assess potential bias caused by lab manipulations, sequencing and downstream bioinformatic processing, a commercially available 16S rRNA mock community, of known composition (Table S1), was included on each plate (Fig. S2) following the manufacturer’s instructions (BEI Resources, USA). The mock community contained DNA of 20 bacterial species (Table S1) in equimolar counts ($10^6$ copies/µL) of 16S rRNA genes. A no-template PCR negative control was also included on each plate, to assess the influence of the PCR reaction, and the efficacy of lab preparation on sequencing (Fig. S2). Four µL of each negative control was mixed with 1 µL of loading dye containing Gel Red (Biotium) and visualized on a 1% agarose gel after 60 minutes at 100 V. None of the negative controls for the DNA extractions nor the PCR reactions contained detectable amounts of DNA prior to submission.

The prepared plates of the Test Phase DNA samples were submitted to Génome Québec (Montréal, Québec) for 16S rRNA amplicon generation and sequencing (Bell *et al*., 2016; Lay *et al*., 2018; Blakney *et al*., 2022). PCR amplification used the S-D-Bact-0341-b-S-17 forward and S-D-Bact-0785-a-A-21 reverse primers, commonly referred to as 341F and 805R, respectively, to generate a 416 bp fragment from the V3-V4 region of the 16S rRNA gene (Klindworth *et al*., 2012). These amplicons were then prepared for paired-end 250 bp sequencing using Illumina’s

MiSeq platform (Génome Québec, Montréal) (Bell *et al.*, 2016; Lay *et al.*, 2018; Blakney *et al.*, 2022). We estimated this should provide a mean of 60 000 reads per sample, which is in line with previous studies that describe bacterial communities (Bell *et al*., 2016; Lay *et al*., 2018; Blakney *et al*., 2022).

<u>Estimating ASVs from MiSeq 16S rRNA gene amplicons</u>

The 16S rRNA gene amplicons generated by Illumina MiSeq were used to estimate the diversity and composition of the bacterial communities present in the bulk soil, seed, rhizosphere and roots from across the Test Phase *B. napus* growth stages. The integrity and totality of the 16S MiSeq data downloaded from Génome Québec was confirmed using their MD5 checksum protocol (Roy *et al*., 2018). Subsequently, all data was managed, and analyzed in R (4.0.3 R Core Team, 2020), and plotted using `ggplot2` (Wickham, 2016).

The `dada2` package (Callahan *et al*., 2016a) was first used to filter and trim all 11 010 728 raw reads, forward and reverse, from the 16S rRNA gene amplicon data generated from the control samples, the mock communities, and the Test Phase *B. napus* samples, using the `filterAndTrim` function (Fig. S2), as described in Blakney *et al*., 2022. Filtered and trimmed reads were then processed through DADA2 for ASV inference (Fig. S3). Default settings were used throughout the DADA2 pipeline, except the DADA inference functions `dadaF` and `dadaR` which used the `pool ='pseudo'` argument, to increase the likelihood of identifying rare taxa. Consequently, the chimera removal function `removeBimeraDenovo` included the `method ='pooled'` argument (Callahan *et al*., 2016b).

ASVs identified from the 16S rRNA gene amplicon data were assigned taxonomy—to the species level where possible—using the Silva database release 138, which adopts the standardized Genome Taxonomy Database taxonomy structure (Yilmaz *et al*., 2013; Parks *et al*., 2018). The quality of the data was assessed using the included controls (Fig. S4); any ASVs identified as chloroplasts, or mitochondria were subsequently removed from the data, as were off-target archaeal ASVs. Rarefaction curves confirmed that we captured the majority of the bacterial communities in both the bulk soil, seed, rhizosphere, and roots (Fig. S5). Test Phase *Brassicaceae* 16S rRNA sequencing data was subsequently re-analysed independently following the described protocol to avoid any biases from the no-template negative controls and the mock communities. These are the Test Phase *B. napus* ASVs which are reported hereafter.

<u>Estimating absolute abundance of bacterial communities by quantitative PCR</u>

To identify any changes in abundance of the bacterial ASVs within the Test Phase *B. napus* growth stages, we estimated the absolute abundance, or size, of the bacterial communities in each Test Phase DNA sample by qPCR (Azarbad *et al*., 2018; Blakney *et al*., 2022). Given the technical-limitations of high-throughput sequencing in assessing abundance, estimating the absolute abundance can provide data to better interpret the bacterial communities (Gloor *et al*., 2017; Props *et al*., 2017; Harrison *et al*, 2020; Jian *et al*., 2020). As such, we quantified the number of 16S rRNA gene sequences present in each DNA sample via qPCR (as Ct values) and then estimated the corresponding community size as the 16S rRNA gene copy number/µg from a standard curve (Zhang *et al*., 2017; Azarbad *et al*., 2018, Fig. S6).

First, a standard curve of 16S rRNA gene copy numbers was constructed. Near full-length 1.5 kb 16S rRNA gene fragments were PCR amplified using the primers PA-27F-YM and PH-R (Bruce *et al*., 1992; Table S2) from DNA extracted from previously used soil samples (Lay *et al*., 2018). The 16S PCR reactions consisted of 11.5 µL $dH_2O$, 5.0 µL of 10X Buffer (Qiagen, Canada), 2.5 µL of 10 µM PA-27F-YM forward and PH-R reverse primers (Alpha DNA, Montréal, Canada;

Klindworth *et al*., 2012), 1.0 μL of dNTPs (Qiagen, Canada), 0.5 μL of *T. aq* polymerase (Qiagen, Canada), and 2 μL of template DNA, for a total volume of 25 μL. PCR amplification was run in an Eppendorf Mastercycle ProS (Germany) thermocycler, and consisted of an initial denaturation of 2 minutes at 95°C, followed by 30 cycles of 30 seconds denaturation at 95°C, 30 seconds annealing at 55°C, and 1 minute elongation at 72°C, before a final elongation of 5 minutes at 72°C (Bell *et al*., 2016; Lay *et al*., 2018; Blakney *et al*., 2022). The amplified 1.5 kb 16S rRNA gene was visualized by a 1% gel electrophoresis, as described above, quantified using the QuBit dsDNA High Sensitivity Kit (Invitrogen, USA), and then serially diluted to $10^{-9}$. One μL of each dilution was then used as template in a 10 μL qPCR reaction.

The 16S rRNA gene qPCR reactions consisted of 5.0 μL of Maxima SYBR Green/ROX qPCR Mix (ThermoFisher Scientific, Canada), 3.4 μL $dH_2O$, 0.3 μL of 10 μM Eub 338 forward and Eub 518 reverse primers (Alpha DNA, Montréal, Canada; Fierer *et al*., 2005). All qPCR reactions were set-up in triplicate in 96-well plates using the Freedom EVO100 robot (Tecan, Switzerland), with a no-template negative control included on each plate. Reactions were run in a ViiA 7 Real-Time PCR System (Life Technologies, Canada) following the same cycling conditions as described previously for the 16S rRNA PCR amplification. The Eub338/Eub518 qPCR reaction amplified a 200 bp region of the V3 region (Muyzer *et al*., 1993; Nogales *et al*., 1999; Bathe & Hausner, 2006; Davis *et al*., 2009). The number of 16S rRNA gene copies present in the serially diluted standard were calculated using the formula (Godornes *et al*., 2007):

$$\text{Number of 16S rRNA gene copies } \mu L^{-1} = \frac{\text{Avogadro's Constant x DNA } (g\ \mu L^{-1})}{\text{Number of base pairs x 600 Daltons}}$$

The standard curve for each diluted sample was plotted, with an $R^2$ value of 0.9938 and an amplification efficiency of -3.2013 (Fig. S6), falling within acceptable values (Fierer *et al*., 2005).

16S rRNA gene copy numbers were then estimated for each Test Phase sample by using 1 μL of a 1:10 dilution of DNA as template in the same 16S rRNA qPCR reaction and cycling conditions as described above for the standard curve. Melt curves generated by 0.5°C increments at the end of the qPCR programme confirmed amplicon specificity, and the 16S rRNA gene copy number was then determined from the standard curve. A correction to determine the absolute abundance of each ASVs inferred from the Test Phase samples was achieved by multiplying the total 16S rRNA gene copy number per μg, as estimated from the qPCR reaction, by the relative abundance matrix of ASVs identified (Azarbad *et al*., 2018; Bakker, 2018).

#### Generating phylogenetic trees

In order to employ phylogeny-based analysis methods, including phylogenetic diversity (PD), and UniFrac distances, we assembled phylogenies for the Test Phase *B. napus* bacterial communities. Following the method described by Callahan *et al*., 2016, 16S rRNA gene sequences for each ASV inferred from the Test Phase data were aligned using a profile-to-profile algorithm (Wang & Dunbrack, 2004) with a dendrogram guide tree using the `decipher` package (Wright, 2016). With the `phangorn` package (Schliep, 2011), the maximum likelihood of each site was calculated using the `dist.ml` function using a JC69 equal base frequency model, before assembling phylogenies using the neighbour-joining method. An optimized GTR nucleotide substitution model was fitted to the phylogeny using the `optim.pml` function. Phylogenies were subsequently added to the `phyloseq` object.

#### α-diversity of the *Brassica napus* bacterial communities

Faith's PD was calculated as an α-diversity index from the *B. napus* samples using the `pd` function from the `picante` package (sum of all branch lengths separating taxa in a community). Log transformed PD indices were confirmed to respect normality. We assessed differences of the mean PD between GS for each SH, and their interactions using a Multi-Factor ANOVA and Tukey's Post-Hoc test for significant groups that respected the assumptions of normality. Normality of the residuals was established with a Shapiro-Wilk test, `shapiro.test`, while the heteroscedascity of residuals was confirmed with using a Bartlett test, `bartlett.test`. For significant ANOVAs, a post-hoc Tukey's Honest Significant Difference test, TukeyHSD, was used to determine which groups were statistically different.

A significant indicator value is obtained if an ASV has a large mean abundance within a group, compared to another group (specificity), and has a presence in most samples of that group (fidelity). The fidelity component complements the differential abundance approach between taxa clusters, which only considers abundance.

β-diversity of the *Brassica napus* bacterial communities

PERMANOVA divides any variation in the ordinated data distance matrix among all the pairs of specified experimental factors. The UniFrac distance index incorporates the phylogenetic relationship of each ASV and their absolute abundance, as estimated by qPCR of the 16S rRNA gene. Determining community distances based only on the number of shared taxa does not account for evolutionary distances between taxa, which are often extremely diverse among microbes. Conversely, using a UniFrac index illustrates how bacterial community composition varies by phylogeny, which provides insight into how different community assembly mechanisms, including dispersal, drift, selection, and speciation, may be at work.

To further characterize what ecological mechanisms may be responsible for changes to the β-diversity, we partitioned it into turnover (i.e. species replacement) and nestedness (i.e. loss/gains that result in poor species richness being a subset of richer sites) components. To take advantage of our abundance data, where turnover and nestedness are analogous to balanced variation & abundance gradients, we first arranged the abundance data between all samples using the `betapart.core.abund` function. Then, to compare each sample to itself through time, we used the `beta.pair.abund` function, with Bray-Curtis dissimilarity distances. As nestedness remained below 0.1 throughout, it was not further analyzed (data not shown). As turnover among compartments was not normally distributed, the non-parametric Kruskal test was used to determine significant, followed by the post-hoc pairwise Wilcox test with an FDR correction. We assessed differences of the mean turnover between GS for each soil history and their interactions as described above for α-diversity, as normality was respected.

### Supplementary Results

As a positive control for our experiment, we included two replicates of the bacterial mock communities, which retained 7430 and 9537 16S rRNA reads (Fig. S4). The mock communities closely resembled each other by ASV composition (Fig. S4), where our pipeline correctly identified all 20 of the bacterial species included in the communities (Fig. S4 & Table S1). These results provide some reassurance that our pipeline ought to be effective in identifying a range of bacterial ASVs present in the experimental samples.

Similar numbers of 16S rRNA reads were retained in the bulk soil samples and rhizosphere samples; means of ~25 429 and 22 467 reads, respectively (Table 1). We identified ~1300 to 1700 ASVs in the bulk soil samples, while the rhizosphere samples had ~800 to 1800 ASVs (Table 1). In contrast, an average of ~3098 16S rRNA reads were retained in the root samples, where only ~100 to 450 ASVs were inferred (Table 1). Finally, we estimated the absolute abundance, or size, of each bacterial community by qPCR amplification of the 16S rRNA gene, where total community sizes ranged from ~200 000 to ~18 000 000 16S rRNA gene copies (Table 1).

### Supplementary Tables

**Table S1**. Bacterial strains included in the mock community (BEI Resources, USA) of known composition, was included on each plate (Fig. S2). The mock community contains DNA of 20 bacterial taxa in equimolar counts ($10^6$ copies/µL) of 16S rRNA genes. Taxa have been provided to illustrate the level of comparison.

| Bacteria | Taxonomy | | |
|---|---|---|---|
| | Phyla | Class | Order/Family |
| *Actinomyces odontolyticus* | Actinobacteriota | Actinobacteria | Actinomycetales (O) |
| *Propionibacterium acnes* | | | Propionibacteriales (O) |
| *Bacteroides vulgatus* | Bacteroidota | | |
| *Deinococcus radiodurans* | Deinococcota | | |
| *Helicobacter pylori* | Campilobacterota | | |
| *Bacillus cereus* | Firmicutes | Bacilli | Bacillales (O)/ Bacillaceae (F) |
| *Listeria monocytogenes* | | | Lactobacillales (O)/ Listeriaceae (F) |
| *Staphylococcus aureus* | | | Staphylococcales (O) |
| *Staphylococcus epidermidis* | | | Staphylococcales (O) |
| *Enterococcus faecalis* | | | Lactobacillales (O)/ Enterococcaceae (F) |
| *Lactobacillus gasseri* | | | Lactobacillales (O)/ Lactobacilliaceae (F) |
| *Streptococcus pneumoniae* | | | Lactobacillales (O)/ Streptococcaceae (F) |
| *Streptococcus agalactiae* | | | |
| *Streptococcus mutans* | | | |
| *Clostridium beijerinckii* | | Clostridia | |
| *Rhodobacter sphaeroides* | Proteobacteria | Alphaproteobacteria | |
| *Neisseria meningitides* | | Gammaproteobacteria | Burkholderiales (O) |
| *Escherichia coli* K12 | | | Enterobacteriales (O) |
| *Acinetobacter baumannii* | | | Pseudomonadales (O)/ Moraxellaceae (F) |
| *Pseudomonas aeruginosa* PAO1-LAC | | | Pseudomonadales (O)/ Pseudomonadaceae (F) |

**Table S2**. Primers used in this study.

| Name | Sequence (5'-3') | Reference |
|---|---|---|
| S-D-Bact-0341-b-S-17 | CCTACGGGNGGCWGCAG | Klindworth *et al*., 2012 |
| S-D-Bact-0785-a-A-21 | GACTACHVGGGTATCTAATCC | Klindworth *et al*., 2012 |
| 16S PA-27F-YM | AGAGTTTGATCCTGGCTCAG | Bruce *et al*., 1992 |
| 16S PH-R | AAGGAGGTGATCCAGCCGCA | Bruce *et al*., 1992 |
| Eub338 | ACTCCTACGGGAGGCAGCAG | Fierer *et al*., 2005 |
| Eub518 | ATTACCGCGGCTGCTGG | Fierer *et al*., 2005 |

**Table S3**. PERMANOVA for all the sampled Test Phase communities identified compartment (bulk soil, rhizosphere, or root), growth stage, and soil history established in the Conditioning Phase, as significant experimental factors. PERMANOVA was calculated using a Bray-Curtis and Weighted Unifrac distance matrix, with 9999 permutations.

| | Bray-Curtis | | | Weighted Unifrac | | |
|---|---|---|---|---|---|---|
| | F Model | $R^2$ | Pr (> F) | F Model | $R^2$ | Pr (> F) |
| Compartment[a] | 2.82430 | 0.05461 | **0.001** | 81.305 | 0.60135 | **0.001** |
| Growth Stage[b] | 1.12762 | 0.02907 | **0.006** | 1.655 | 0.01632 | 0.090 |
| Soil History[c] | 1.14988 | 0.01482 | **0.016** | 2.924 | 0.01442 | **0.025** |
| Compartment ~ Growth Stage | 1.29513 | 0.05008 | **0.001** | 2.576 | 0.03811 | **0.005** |
| Compartment ~ Soil History | 1.03763 | 0.02675 | 0.154 | 1.458 | 0.01438 | 0.156 |
| Soil History ~ Growth Stage | 0.97479 | 0.05026 | 0.762 | 0.767 | 0.01514 | 0.724 |
| Compartment ~ Soil History ~ Growth Stage | 1.01258 | 0.07832 | 0.268 | 1.150 | 0.03402 | 0.296 |

a, Rhizosphere or roots
b, Seed, seedling, rosette, bolting, or flower
c, Monocrop canola, wheat-canola rotation, or pea-barely-canola rotation

### Supplementary Figures

**Fig. S1**

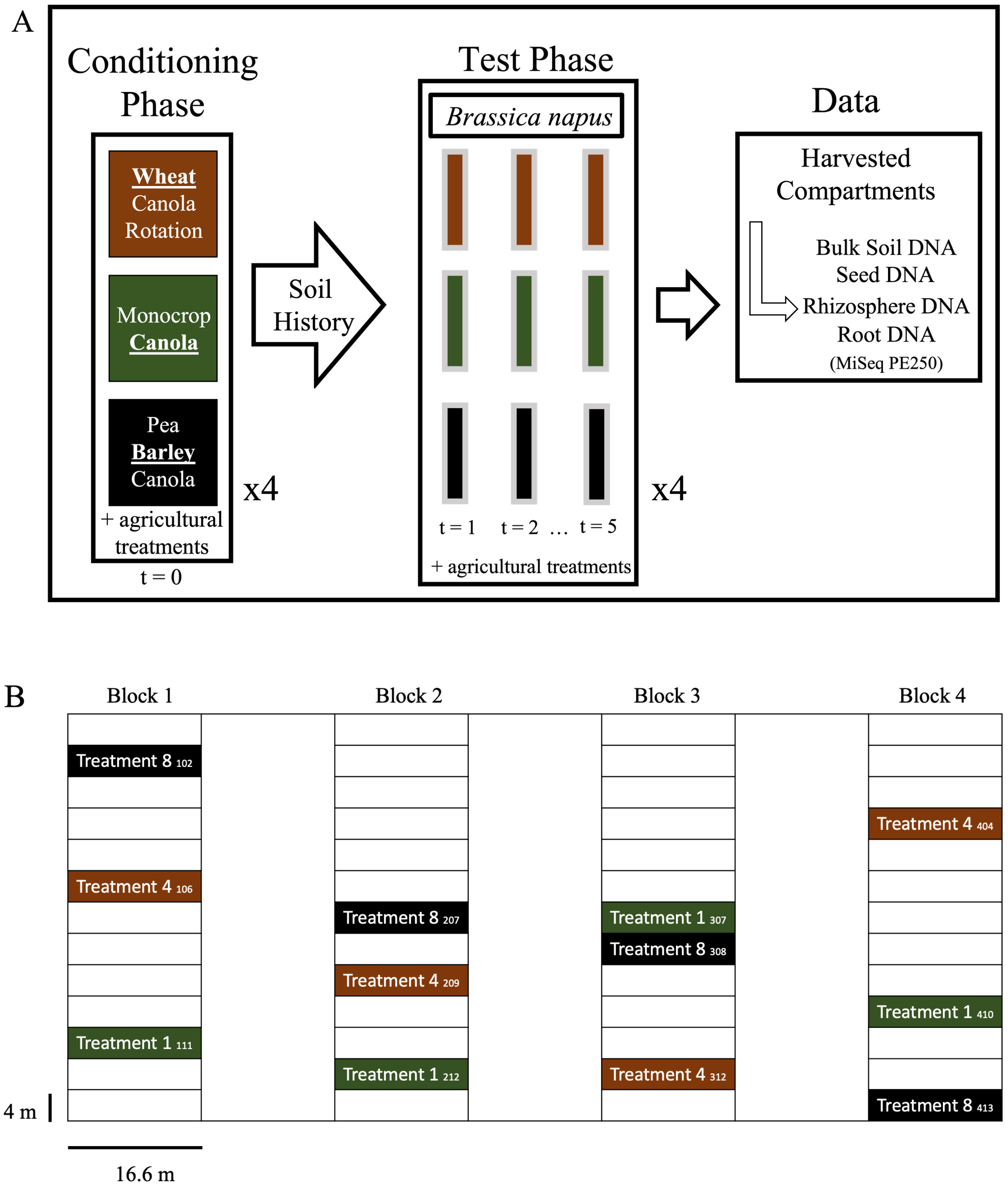

**Figure S1**. Conceptual design of the experiment. (A) Soil history was established during the growing season of 2018 (Conditioning Phase, t = 0), while the effect of soil history on *Brassica napus* bacterial rhizosphere and root communities, or associated bulk soil, was observed the following season, in 2019 (Test Phase, t = 1, 2, 3 … 5). Samples were harvested throughout the growing season at five growth stages; seed, seedling, rosette, bolting, and flower. (B) Field plan for the experiment. The experimental design was a split-plot replicated in four complete blocks. In the 'Conditioning Phase', three soil history treatments were randomly assigned: treatment 1 (green) another generation of canola (*B. napus* L., cv. L252LL) as a monocrop, treatment 4 (brown) spring wheat (*Triticum aestivum* cv. AAC Brandon), as the wheat phase of a two-year crop rotation with *B. napus*, and treatment 8 (black) barley (Hordeum vulgare cv. Canmore), as a three-year crop rotation with *B. napus* and pea (*Pisum sativum* L. cv AAC Lacombe). Plot numbers appear in subscript.

547 **Fig. S2**

548

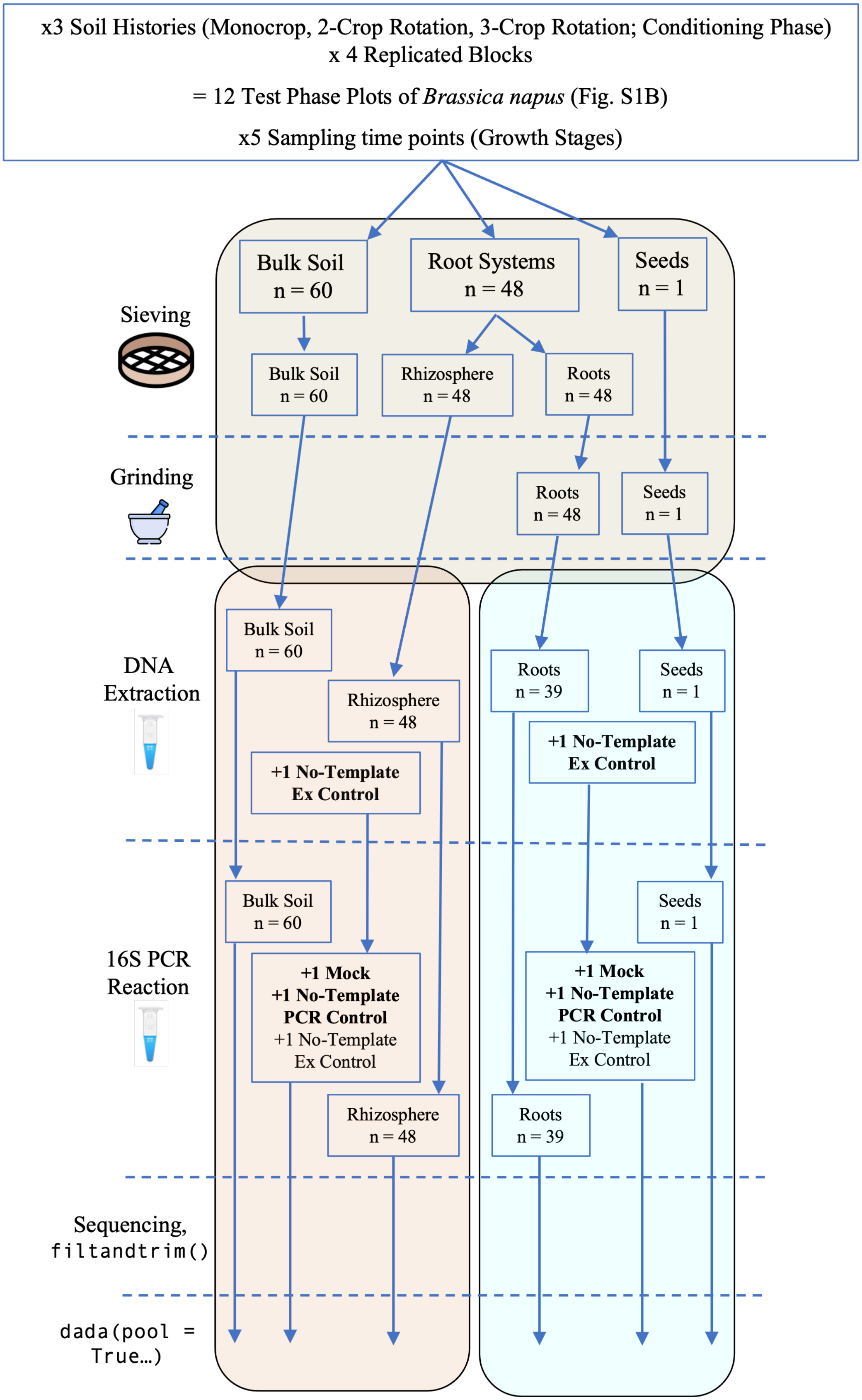


549

**Figure S2**. Organization of our lab workflow for the Test Phase *Brassicaceae* samples from harvest to generating amplicon sequence variants (ASVs). The Test Phase *Brassicaceae* samples were harvested in from 12 plots organized as split plots that were randomly assigned three soil history treatments (Fig S1B). Each plot was harvest at five growth stages (seed, seedling, rosette, bolting and flower); note the seed was all the same for each plot (n = 1). Bulk soil samples were taken from each plot at each growth stage (n = 60). In the field, each plant had its root systems and associated immediately flash-frozen in liquid nitrogen and kept on ice. In the lab, samples were sieved to remove debris (rocks, undecomposed straw, twigs etc…), and root systems were divided into rhizosphere and root samples (n = 48 each). Roots and seeds were then ground in liquid nitrogen, and DNA was extracted from all the Test Phase samples. No-template extraction controls were included to assess what contaminates, or biases, the extraction kits (Machery-Nagel Nucleospin Soil gDNA kit, at left in brown, and Qiagen Plant DNEasy kit in green) might impart. We also included no-template PCR negative controls, and confirmed by gel electrophoresis that none of the no-template extraction controls, nor the no-template PCR controls, contained DNA. To help identify sequencing biases, or bath effects, a replicate of the bacterial mock community (BEI Resources, USA) was included on each plate submitted for sequencing. All DNA samples were submitted to Génome Québec for 16S rRNA PCR amplification, library preparation, and paired-end 250 bp Illumina MiSeq sequencing. All reads were subsequently trimmed and processed through the DADA2 pipeline for ASV inference.

**Fig. S3**

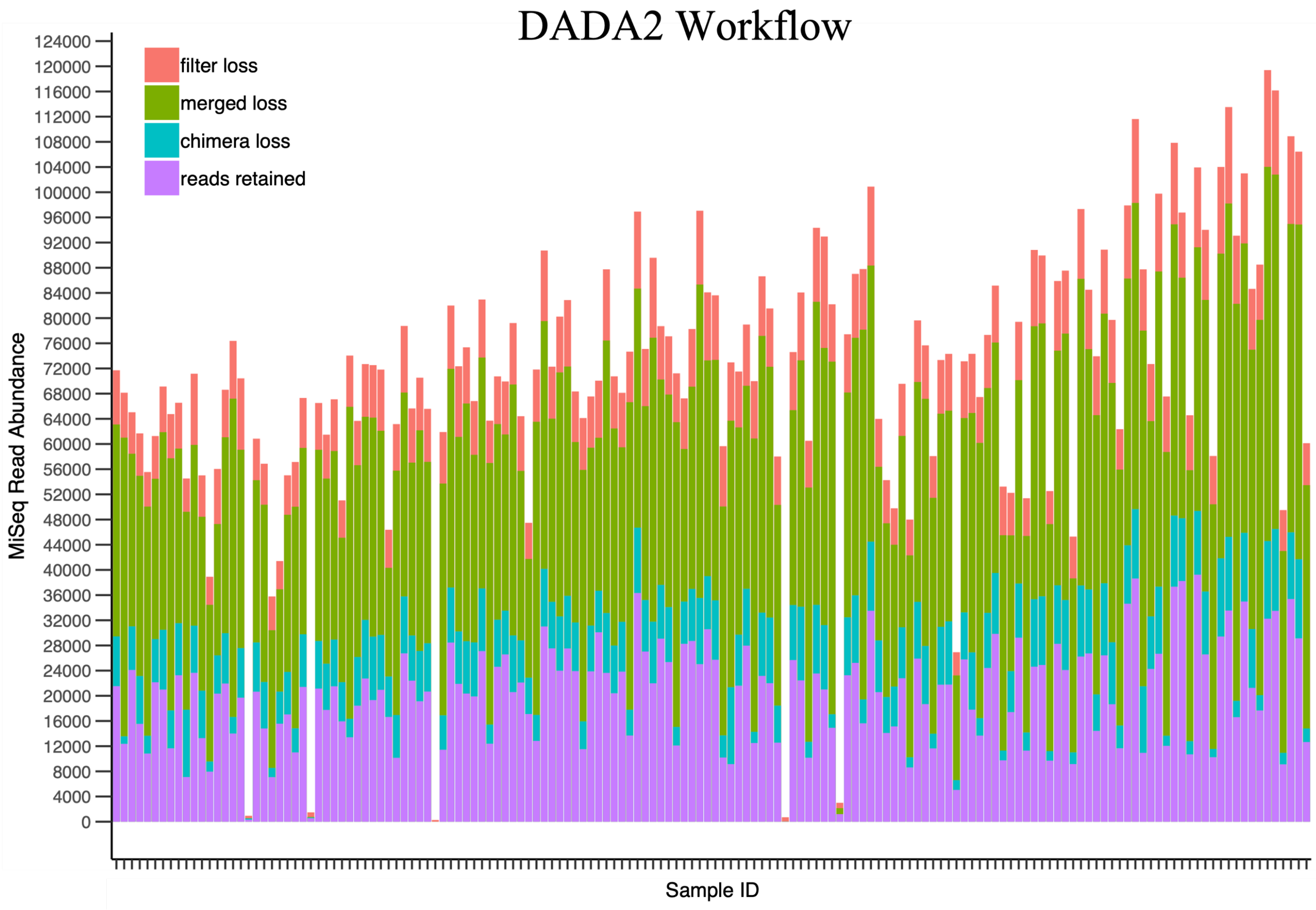


**Figure S3**. The DADA2 workflow processed 11 010 728 raw reads, produced from one lane of sequencing via Illumina's MiSeq at Génome Québec, and retained 2 770 390 reads which were used to infer amplicon sequence variants (ASVs).

**Fig. S4**

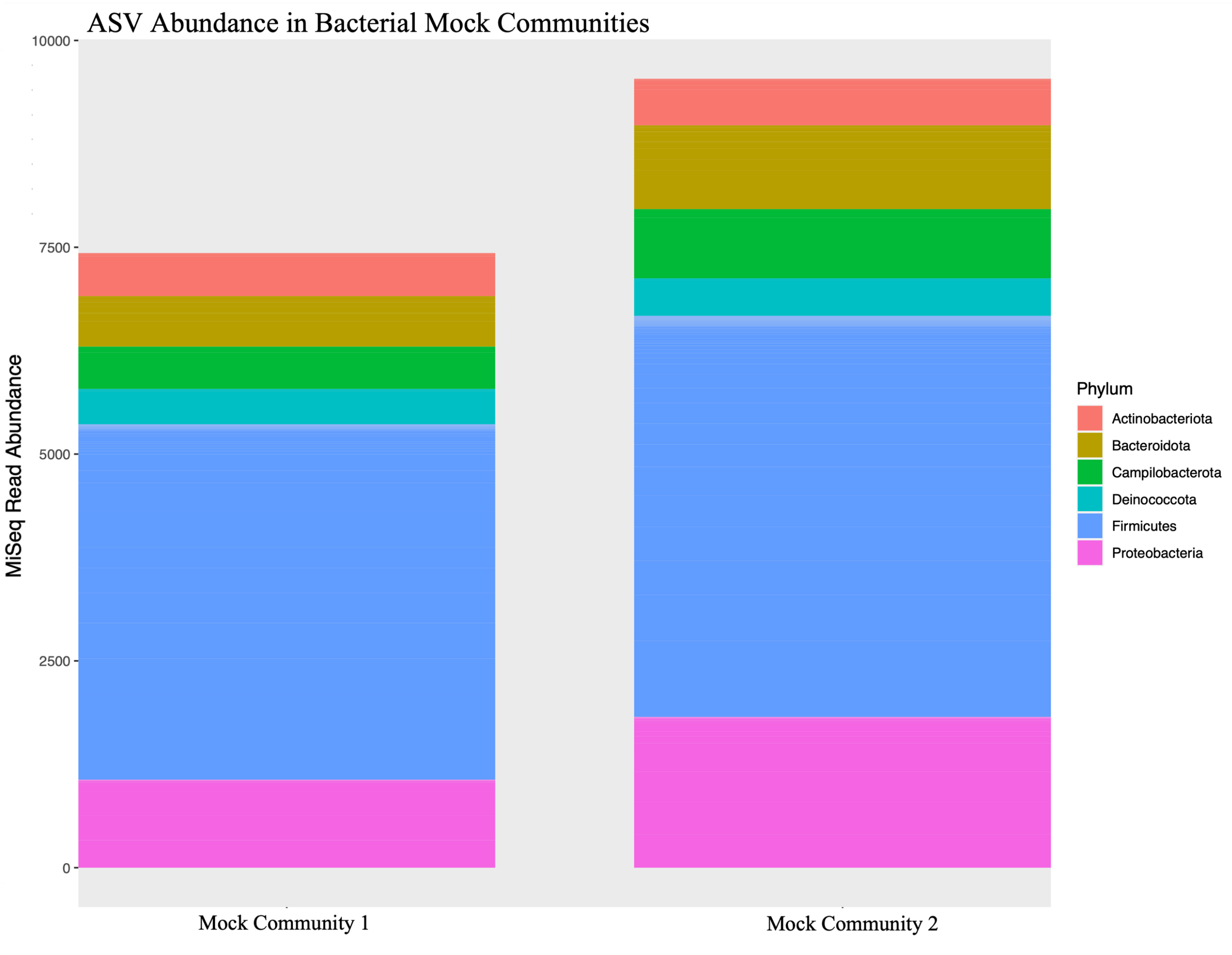


**Figure S4**. High-quality MiSeq reads retained through the DADA2 pipeline from among the mock community replicates, were inferred as inferred amplicon sequence variants (ASVs), and assigned taxonomy using the Silva database, represented here as phlya. Every bacterial group included in the mock community was detected in both replicates, and no others. The *Actinobacteria*, and *Deinococcus*, were the most accurate between what was included in the community, and retrieved in our pipeline. Community 2 showed an expansion among the ASVs identified as *Bacteriodetes*, *Campilobacterota*, *Firmicutes*, and *Proteobacteria*.

**Fig. S5**

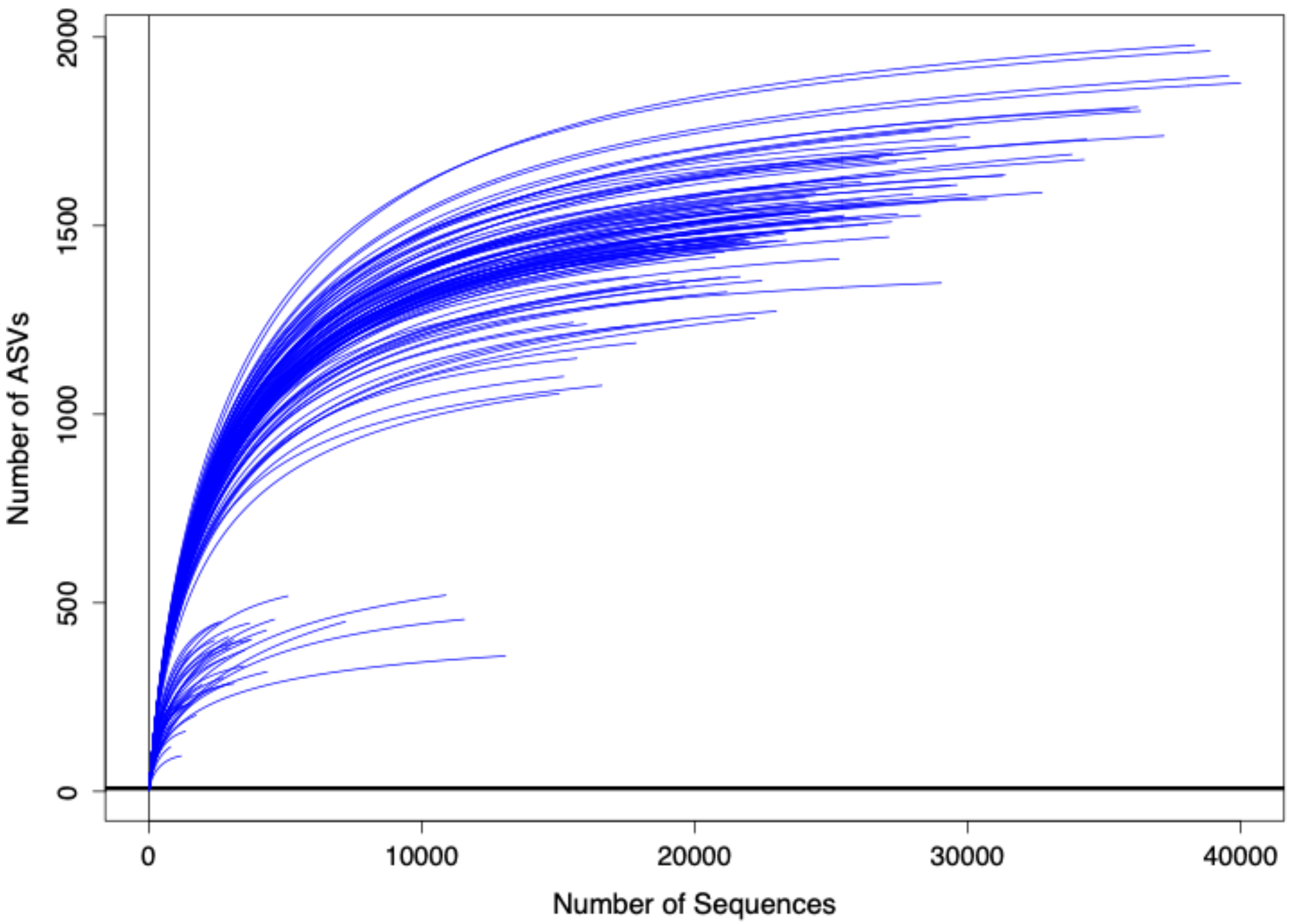


**Figure S5**. Rarefaction curves illustrated that the majority of the bacterial communities were identified in the samples harvested in 2019 from the bulk soil, rhizosphere, and roots, from five growth stages during the Test Phase of a multi-year crop rotation, in Lacombe, Alberta.

**Fig. S6**

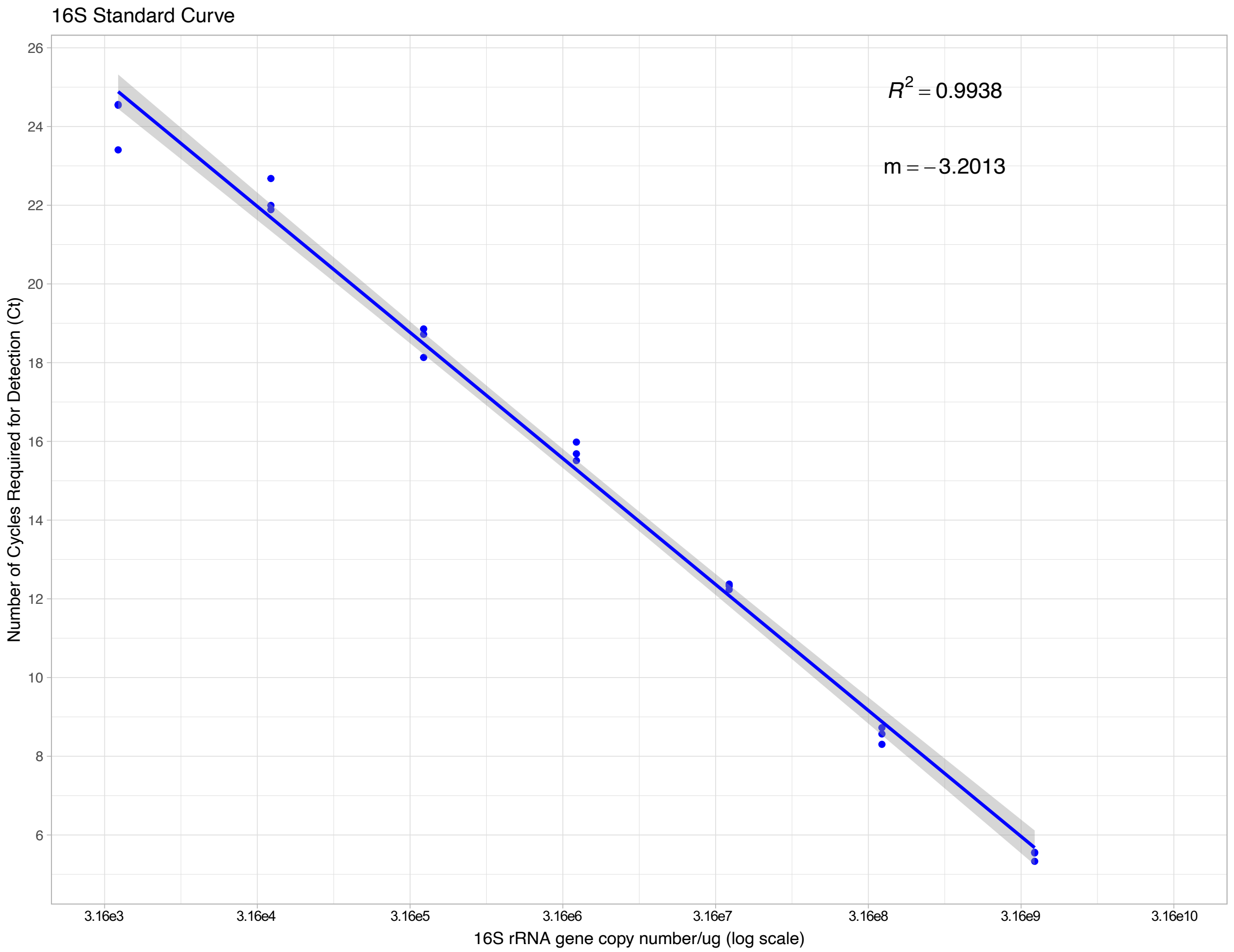


**Figure S6**. A standard curve of the 16S rRNA gene copy numbers (X-axis) versus the number of cycles required for detection (cycle threshold, Ct, Y-axis), as determined from the serial dilution of a quantified 16S rRNA gene.

**Fig. S7**

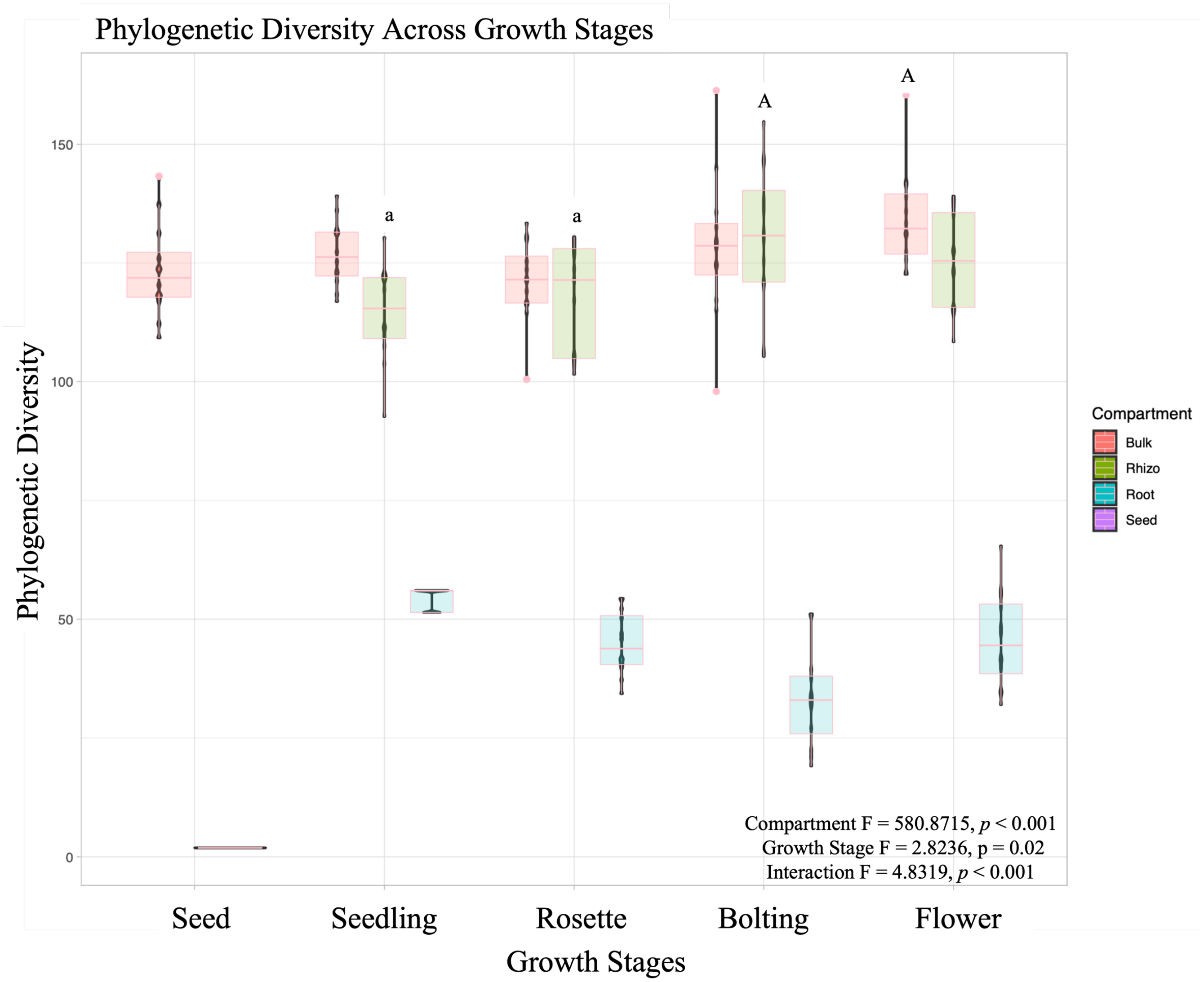
Phylogenetic Diversity Across Growth Stages
Phylogenetic Diversity
150
100
50
0
a
a
A
A
Compartment
Bulk
Rhizo
Root
Seed
Compartment F = 580.8715, *p* < 0.001
Growth Stage F = 2.8236, p = 0.02
Interaction F = 4.8319, *p* < 0.001
Seed
Seedling
Rosette
Bolting
Flower
Growth Stages

**Figure S7**. Phylogenetic diversity was significantly different between compartments ($p < 0.001$) in bacterial communities throughout the *Brassica napus* growing season. Root and seed communities were significantly different from each compartment across growth stages. Bulk soil and rhizosphere communities were strikingly similar across growth stages, except for the flower and bolting stage, respectively, which were both more diverse than the rhizosphere communities at the seedling and rosette stages. Diversity across growth stages was tested with a Multi-Factor ANOVA, which confirmed that the compartments and the Test Phase *B*. *napus* growth stages were significant and did interact. Statistically significant groups were identified using Tukey's post-hoc test.

**Fig. S8**

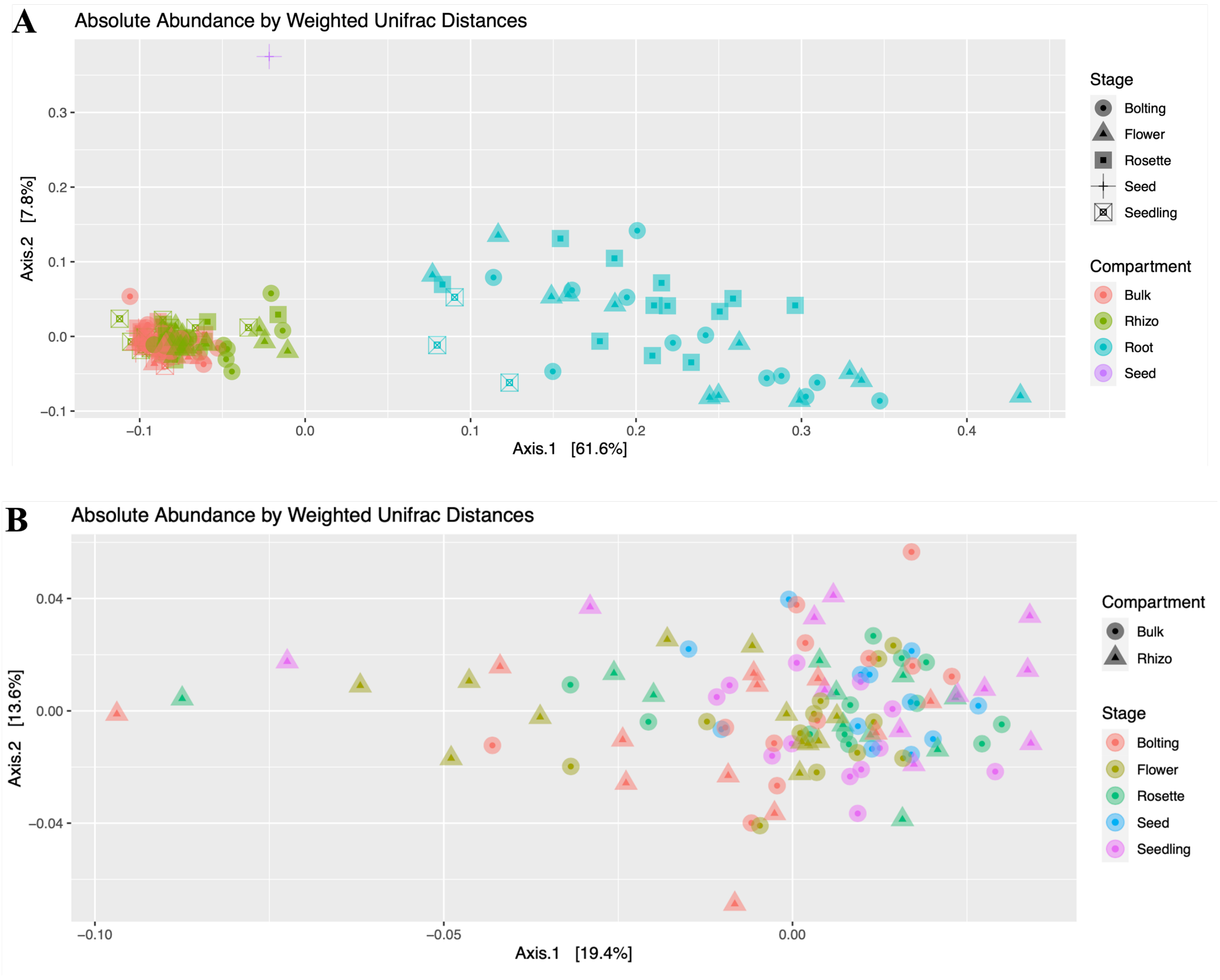
A
Absolute Abundance by Weighted Unifrac Distances
Axis.2 [7.8%]
Axis.1 [61.6%]
0.3
0.2
0.1
0.0
−0.1
−0.1
0.0
0.1
0.2
0.3
0.4
Stage
Bolting
Flower
Rosette
Seed
Seedling
Compartment
Bulk
Rhizo
Root
Seed
B
Absolute Abundance by Weighted Unifrac Distances
Axis.2 [13.6%]
Axis.1 [19.4%]
0.04
0.00
−0.04
−0.10
−0.05
0.00
Compartment
Bulk
Rhizo
Stage
Bolting
Flower
Rosette
Seed
Seedling

**Figure S8**. Bulk soil and rhizosphere bacterial communities were more phylogenetically similar than to root or seed communities. (A) Principal co-ordinate analysis illustrated that seed and root bacterial communities were phylogenetically distinct from bulk soil and rhizosphere communities. Bacterial root communities were more similar at the seedling and rosette stages, and appeared more diverse over time. Axis 1 and 2 captured 69.4% of the variability among the bacterial communities. (B) Bacterial communities from the bulk soil and rhizosphere were more phylogenetically similar at the seed and seedling stages, and increasingly diverse at following growth stages. Axis 1 and 2 captured 35% of the variability among the bacterial bulk soil and rhizosphere communities. Principal co-ordinate analysis were plotted using UniFrac distances weighted by absolute abundance, where phylogenetically similar communities were plotted closer together.

**Fig. S9**

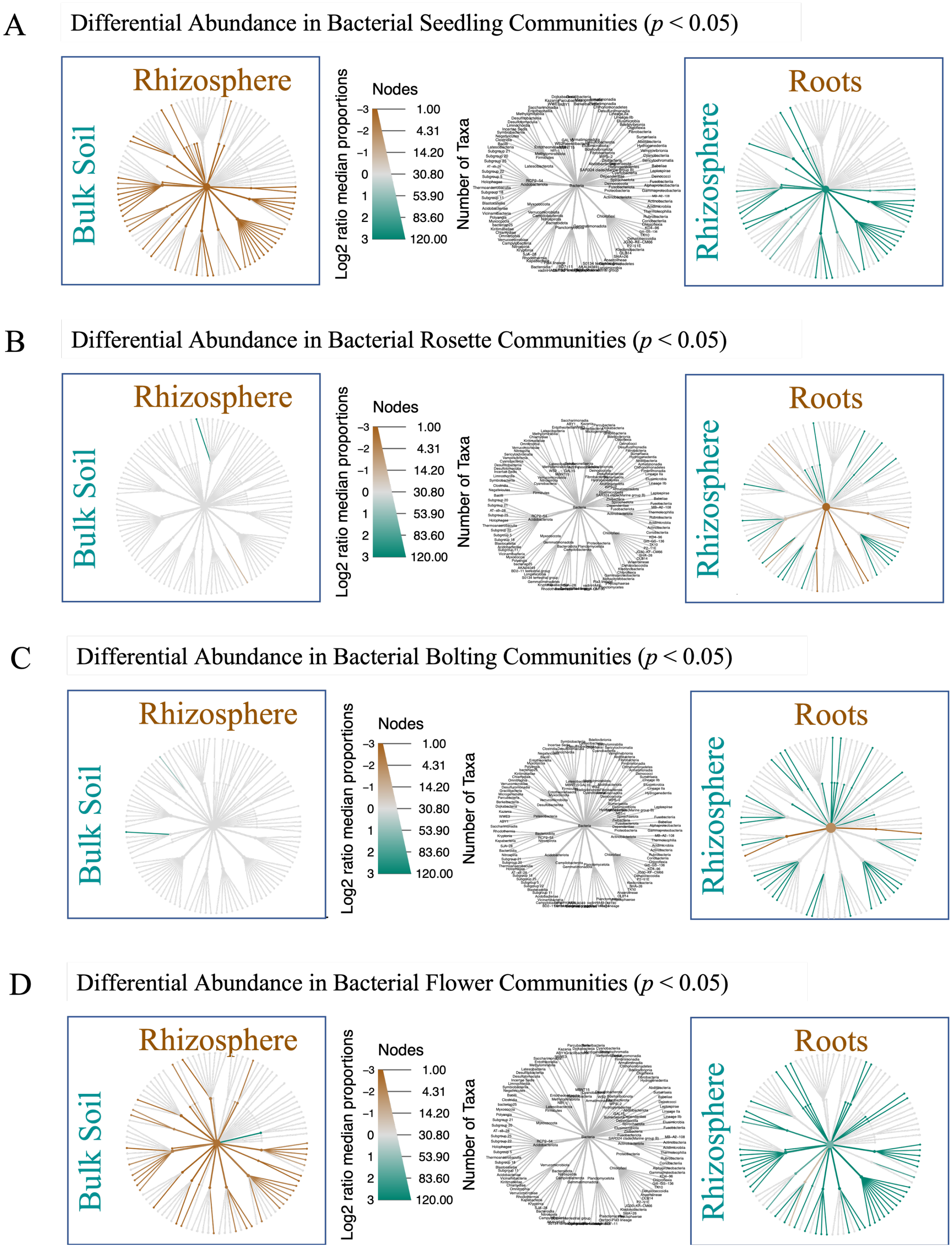
A
Differential Abundance in Bacterial Seedling Communities ($p < 0.05$)
B
Differential Abundance in Bacterial Rosette Communities ($p < 0.05$)
C
Differential Abundance in Bacterial Bolting Communities ($p < 0.05$)
D
Differential Abundance in Bacterial Flower Communities ($p < 0.05$)
Rhizosphere
Bulk Soil
Roots
Rhizosphere
Nodes
Log2 ratio median proportions
Number of Taxa
−3 −2 −1 0 1 2 3
1.00 4.31 14.20 30.80 53.90 83.60 120.00

**Figure S9**. Specific bacterial taxa were significantly different between adjacent compartments at specific *Brassica napus* growth stages. (A) Bacterial rhizosphere communities at the seedling stage were enriched in taxa compared to the bulk soil and root communities. (B) At the rosette stage, bulk soil and rhizosphere communities were relatively similar, while both the rhizosphere and root communities were enriched in different taxa. (C) The bacterial communities in the bulk soil and rhizosphere were also similar at the bolting stage, though the the rhizosphere and root communities both exhibited specific enrichments in different taxa. (D) Bacterial rhizosphere communities were enriched in taxa compared to both the bulk soil and root communities. The abundance of each taxonomic group was compared between compartments, using the using the non-parametric Kruskal test and the post-hoc pairwise Wilcox test, with the FDR correction. Taxa that were significantly (*p*. adj $< 0.05$) more abundant were highlighted.
